# PEPITEM Tripeptides and Peptidomimetics: Next-Generation Modulators of Inflammation in Immune-Mediated Conditions

**DOI:** 10.1101/2024.10.14.618151

**Authors:** Anella Saviano, Bonita Apta, Samantha Tull, Laleh Pezhman, Areeba Fatima, Antonio Mete, Mustafa Sevim, Myriam Chimen, Anna Schettino, Noemi Marigliano, Helen M. McGettrick, Asif J. Iqbal, Francesco Maione, G. Ed Rainger

**Affiliations:** ImmunoPharmaLab, Department of Pharmacy, School of Medicine and Surgery, University of Naples Federico II, Via Domenico Montesano 49, 80131, Naples, Italy; Department of Cardiovascular Sciences, University of Birmingham, Birmingham, B15 2TT, UK; Department of Inflammation and Ageing, University of Birmingham, Birmingham, B15 2TT, UK; Medsyndesign Ltd, Advanced Technology Innovation Centre, 5 Oakwood Drive, Loughborough, LE11 3QF, UK

**Author notes:** Authors contributed equally to this work (Anella Saviano, Bonita Apta, Samantha Tull). Authors contributed equally to this work (Helen M. McGettrick, Asif J. Iqbal, Francesco Maione, G. Ed Rainger). **Correspondence:** Professor Ed Rainger, Department of Cardiovascular Sciences, The Medical School, University of Birmingham, Birmingham, B15 2TT, UK.; Professor Francesco Maione, Head of *ImmunoPharmaLab*, Department of Pharmacy, School of Medicine and Surgery, University of Naples Federico II, Via Domenico Montesano 49, 80131, Naples, Italy.

## Abstract

PEPITEM is an immune-modulatory peptide that effectively regulates inflammation and mitigates immune-mediated inflammatory diseases (IMIDs). Here we identify two independently active tripeptide pharmacophores within PEPITEM and engineered peptidomimetics with enhanced pharmacodynamic properties. These peptidomimetics regulate T-cell trafficking *in vitro* and reduce T-cell, neutrophil and macrophage numbers in the inflamed peritoneal cavity i*n vivo*. In a plaque psoriasis model, topical administration reduced disease severity, inflammation and immune cell infiltration, while regulating cytokine release in macrophages and fibroblasts, as well as keratinocyte proliferation. Th1 and Th17 cell abundance, along with their cytokines, was reduced in secondary lymphoid organs. This expanded functional repertoire of PEPITEM and its derivatives provides innovative tools for countering immune and stromal cell-induced pathology in IMIDs, paving the way for a novel class of anti-inflammatory agents.

## 1 Introduction

Despite substantial therapeutic innovation over the past two decades, immune mediated inflammatory diseases (IMIDs) represent an area of inadequately controlled morbidity and mortality. These disorders, which include common autoimmune and chronic inflammatory diseases such as rheumatoid arthritis (RA), inflammatory bowel disease (IBD), lupus and psoriasis (and many others) require a range of immunomodulatory agents from chemical drugs through to synthetic biologic agents as standard of care therapies across the disease spectrum^1^. Although this results in polypharmacy for many patients, there remains significant unmet need. Biologics, and many of the first line therapies used for less severe disease are disadvantaged by serious side effects (e.g. serious infections, and malignancies), often associated with systemic immune suppression, which is a major drawback of targeting pro-inflammatory mediators and their downstream pathways^2^. Moreover, there is no curative treatment to restore full control of the immune system, and currently, there are no therapeutics which rebuild, or reverse tissue damage caused by inflammation. Response rates to therapy are not durable. For example, anti-tumor necrosis factor (TNF) therapy in RA is effective in approximately 60% of patients and this is reduced to 20% after 2 years due to primary non-response, secondary non-response or intolerance^3^. Moreover, few therapies achieve disease remission, which has an impact on patient adherence to prescribed treatments. This also means that current therapeutic approaches universally fail to impose control over homeostatic processes required for tissue regeneration and subsequent resolution of the inflammatory insult.

Chronic inflammation is an underlying causative factor in the pathology of all IMIDs. However, inflammation is ordinarily an acute, tightly regulated physiological response to injury, trauma or infection that is necessary to heal damaged tissues. The desired outcomes of the inflammatory response are thus sterilisation (if required), removal of the initiating trigger and repair of the affected tissues and organs, with ideally, their return to homeostatic function. As part of this process, we now know that there are specific pro-resolution pathways that have evolved to achieve this, for example specialised pro-resolving lipid mediators such as the protectins and resolvins and the induction of the anti-inflammatory cytokine interleukin (IL)-10 ^4,5^. Moreover, we now appreciate that chronic inflammation does not only represent a prolonged over activity of pro-inflammatory mediators at the site of disease, it also encompasses loss of function of pro-resolution pathways. An example of just such a scenario is the PEPITEM (<u>PEP</u>tide Inhibitor of <u>T</u>rans-<u>E</u>ndothelial <u>M</u>igration) pathway^6^. PEPITEM is a 14 amino acid (AA) peptide (SVTEQGAELSNEER), proteolytically excised from 14.3.3. zeta/delta (14.3.3. ζδ) protein and secreted by B-lymphocytes during an inflammatory response ^6^. PEPITEM has powerful regulatory functions in the inflammatory response, where it operates to limit the number of pro-inflammatory effector T-cells and macrophages that traffic to tissues under inflammatory conditions ^6, 7, 8, 9^. The PEPITEM pathway undergoes a process of functional decline during human ageing ^6^, possibly associated with lifestyle choices such as obesity ^7^, and this is strongly association with the acquisition of chronic diseases (IMIDs) such as type-1 diabetes and RA ^6^. Studies in animal models of disease show that provision of exogenous peptide by injection can reduce inflammation and disease scores in Sjogren’s Syndrome ^6^, uveitis ^6^, ischaemic disease ^6^; bacterial infectious disease [salmonella infection] ^6^ kidney disease associated with systemic lupus erythematosus (SLE) ^8^, obesity ^7^ and ageing ^9^. Thus, it is possible that the loss of PEPITEM function and its homeostatic activity in the inflammatory response is common to many IMIDs, and its replacement at an early stage of the disease process could alter the progression and severity of symptoms across a wide range of diseases with differing aetiologies.

The use of peptides such as PEPITEM for therapeutic intervention in disease can be problematic. The cut off for renal clearance of plasma borne molecules is in the order of 50kDa, resulting in rapid clearance of small peptides (e.g., PEPITEM-MW 1.549kDa) from the blood by the kidneys ^10^. Moreover, many peptides are subject to enzymatic degradation by circulating peptidases ^11^. To overcome such barriers, peptides can be modified to increase molecular weight or alter resistance to peptidase attack. These changes are part of a wider strategy to improve the characteristics of absorption, distribution, metabolism and excretion (ADME) to optimise the pharmacokinetics of the peptide ^12^. Here we undertook an optimisation programme to improve the pharmacokinetics of PEPITEM and we tested new pharmacophores and peptidomimetics using *in vitro* and *in vivo* models of inflammation. To model efficacy in an IMID we have used topical application of PEPITEM to the skin to intervene in a model of plaque psoriasis. Psoriasis is an autoimmune disease of the skin in which environmental triggers initiate hyperplastic proliferation of cells in the epidermis ^13^. These cells release powerful inflammatory mediators such as TNF-α, IL-1β, with the IL-23/IL-17 axis thought to play a dominant role in disease progression and severity ^13^. Disease can be managed using topical and/or systemic treatments that are anti-inflammatory, antiproliferative or immune suppressive, with the limitations due to side effects already outlined above ^13^. Here we show that topical application of PEPITEM, its pharmacophores and the peptidomimetics derived from the parent sequences have powerful disease modifying actions in psoriatic disease.

## 2. Materials and methods

### 2.1 Design and synthesis of peptides and peptide mimetics

Peptides sequences identified in house, or peptidomimetics designed by Medsyndesign Ltd, Loughborough, UK) were synthesised using standard solid peptide synthesis chemistry ^14^ and purified >95% by HPLC by Biosynth Laboratories Ltd. (Billingham, UK; certificate of analysis and typical HPLC trace for PEPITEM can be found in **Supplementary Fig. 1**).

### 2.2 Cell culture

Human dermal blood endothelial cells (HDBEC; Lot #423Z004, Promocell) were stored on liquid nitrogen at passage 4. Prior to experiment they were thawed and grown in low serum (2% v/w) endothelial cell growth medium MV (PromoCell; C-22120), supplemented with 10% fetal calf serum (FCS; PromoCell; C-37310), endothelial cell growth extracted from hypothalamic (ECGS/H; Promocell; C-30120), in 25 cm^2^ culture flasks until confluent, and then sub-cultured into 24-well plates (2×10^5^ cells/well) and cultured overnight at 37 °C, 5% CO_2_ prior to static adhesion assay ^6^. Murine macrophage J774A.1 (ATCC® TIB-67), mouse embryonic fibroblast (NIH-3T3, ATCC® CRL-1658) and human epidermal keratinocyte (HaCaT, ATCC® PCS-200-011) cell lines were cultured in 100×20 mm dishes (5×10^5^ cells/dish) in Dulbecco’s modified Eagle’s medium (DMEM; Corning; 10-013-CV) supplemented with 10% fetal bovine serum (FBS; Sial; YourSIAL-FBS-SA), 100 U mL^−1^ penicillin, 100 μg mL^−1^ streptomycin (Corning; 30-002-CI), and 25 mM (4-(2-hydroxyethyl)-1-piperazineethanesulfonic acid (HEPES; Sigma-Aldrich; H0887) in a humidified 5% carbon dioxide atmosphere at 37 °C ^15, 16,, 17^.

### 2.3 Static adhesion assay

Ethical approval for this project was awarded by the University of Birmingham. Local Ethical Review Committee (ERN_12-0079). Healthy volunteers recruited for this study were required to be >18 years old, to not have any active infections or immune-related diseases, and not to have had any vaccines <3 month prior to sample collection. Human blood from (23-60 years) was collected into ethylenediamino tetraacetic acid (EDTA)-coated vacuettes. Peripheral blood mononuclear cells (PBMC)s were isolated from blood using a histopaque (Sigma-Aldrich) gradient, as previously described ^6^. The PBMC pellet was resuspended in medium 199 (M199; Gibco) with 0.15% bovine serum albumin (BSA; Sigma-Aldrich) or in MACS buffer (Miltenyi Biotech, UK). Monocytes were removed by panning on culture plastic to yield peripheral blood lymphocytes (PBL) and adjusted to 1×10^6^ cells/mL prior to the static adhesion assay ^6^.

HDBEC were stimulated with 100 U mL^−1^ TNF-α (R&D Systems, UK) and 10 ng mL^−1^ interferon-gamma (IFNγ; PeproTech, UK) for 24 h prior to washing and addition of PBL. PBL were either untreated or treated with different concentrations of PEPITEM, its tripeptide pharmacophores or their peptidomimetics during the assay. PBL were permitted to migrate for 20 min (37 °C, 5% CO_2_). Non-adherent PBL were then removed by gentle washing with M199 supplemented with 0.15% BSA. Wells were imaged using a phase-contrast Olympus IX70 microscope (Olympus, UK) and digitised images analysed off-line using Image-Pro 6.2 software (Media Cybernetics; Maryland, USA) as previously described^6^. Total PBL adhesion was expressed as cells per mm^2^ per number of PBL added. PBL transendothelial migration was calculated as a percentage of the total number of adhered cells ^6^

### 2.4 Cell viability assay and treatments

MTT assay was conducted to determine the cytotoxicity of PEPITEM, its tripeptide pharmacophores or their peptidomimetics on J774A.1 macrophage cell line ^15^. Briefly, J774A.1 cells (5×10^3^ per well) were seeded in 96-well plates and, after overnight incubation, were treated with PEPITEM, control peptide (the 14aa proximal to the PEPITEM sequence; EQAERYDDMAACMK), SVT-acid, SVT[NH-Ethyl], SVT-amide, AC-QGA-acid or [pGlu]GA-amide at different concentrations (1, 3, 10, 20 and 30 ng mL^−1^). After 4 and 24 h, 10 µL of MTT (BioBasic; T0793) solution (5 mg mL^−1^ in phosphate-buffered saline, PBS; pH 7.4) was added to each well, and the plates were incubated for 3 h at 37 °C in the dark. Then, the medium was removed, and the obtained formazan crystals were dissolved in 150 µL of dymethyl sulfoxide (DMSO) for 15 min. The spectrophotometric absorbance was measured using a microtiter enzyme-linked immunosorbent assay reader (Multiskan^™^ GO Microplate Spectrophotometer; Thermo Scientific^™^) at 540 nm. The percentage of cell viability was determined by the following formula: OD of treated cells/OD of control ×100. For further *in vitro* experiments, following 70% confluency, J774A.1 macrophage and NIH-3T3 fibroblast cells were pre-treated for 15 min with the aforementioned compounds at the concentration of 10 ng mL^−1^ (highest non-cytotoxic concentration) before being stimulated with lipopolysaccharide from *Escherichia coli-*O111:B4 (LPS, 10 µg mL^−1^, Product Number: L2630; Sigma-Aldrich). for 24 h. Supernatants were collected and stored at −80 °C for IL-6 and TNF-α enzyme-linked immunosorbent assay (ELISA) analysis. Protein concentration was determined by the Bio-Rad protein assay (Bio-Rad, Milan, Italy) as previously described ^15, 16^.

### 2.5 Cell proliferation assay

Psoriasis-like keratinocytes model was established by stimulating HaCaT cells with M5 cytokines cocktail consisting of 2.5 ng mL^−1^ of each TNF-α (210-TA/CF), Oncostatin M (295-OM/CF), IL-1α (200_LA/CF), IL-17 (317-ILB) and IL-22 (782-IL/CF, all from R&D System, Italy) as previously described ^17, 18^. Briefly, human keratinocytes (5×10^3^ per well) were seeded in 96-well plates and, after allowed to adhere overnight, were pre-treated for 15 min with PEPITEM, control peptide, SVT-acid, SVT[NH-Ethyl], SVT-amide, AC-QGA-acid or [pGlu]-GA-amide at the concentration of 10 ng mL^−1^ before stimulation with M5 mix. After 72 h, cell proliferation was assessed by using an MTT assay as described above ^19^.

### 2.6 ELISA assay

The levels of IL-6 (DY406; R&D System) and TNF-α (860.040.192; Diaclone) in J774A.1 and NIH-3T3 supernatants were measured at 24 h using commercially available ELISA kit ^20^. Briefly, 100 μL of supernatants, diluted standards, quality controls, and dilution buffer (blank) were applied on a pre-coated plate with the monoclonal antibody for 2 h. After washing, 100 μL of biotin-labelled antibody was added, and incubation continued for 1 h. The plate was washed, and 100 μL of the streptavidine-HRP conjugate or enzyme conjugate was added, and the plate was incubated for a further 30 min period in the dark. The addition of 100 μL of the substrate and stop solution represented the last steps before the reading of absorbance (measured at 450 nm) on a microplate reader (Multiskan™ GO Microplate Spectrophotometer; Thermo Scientific™). Antigen levels in the samples were determined using a standard curve, normalized to supernatant levels, and expressed as pg mL^−1^ ^21, 22^.

### 2.7 Animals

Studies in the UK (*in vivo* model of peritonitis) were conducted in accordance with UK Home Office regulations with appropriate ethics. Six to eight-week-old C57Bl/6 J wild type (WT) mice (male) from Charles River were housed at the University of Birmingham animal unit with free access to food (rodent 5LF2 diet; IPS LabDiet) and water under SPF environmental conditions at 21 ± 2 °C, 55 ± 10% relative humidity and a 12 h light-dark cycle. Conditions were qualified by screening using Federation of European Laboratory Animal Science Associations approved methods.

For studies in Italy (*in vivo* model of psoriasis), all animal care and experimental procedures complied with reporting of *in vivo* experiments (ARRIVE) guidelines, international and national law and policies and were approved (Authorisation number: 507/2022-PR) by the Italian Ministry of Health (EU Directive 2010/63/EU for animal experiments, and the Basel declaration including the 3Rs concept). Female, 8–12-week-old BALB/c were purchased from Charles River (Milan, Italy). Animals were housed under specific pathogen-free conditions in ventilated cages under controlled temperature and humidity, on a 12 h light/dark cycle and allowed *ad libitum* access to standard laboratory chow diet and sterile water from Charles River (Milan, Italy). All procedures were carried out to minimize the number of animals used (n=6 per group) and their suffering. Experimental study groups were randomized, and their assessments were carried out by researchers blinded to the treatment groups.

### 2.8 Induction of zymosan-induced peritonitis

For the *in vivo* model of peritonitis, inflammation was induced by intraperitoneal (IP) injection of 0.1 mg zymosan A (Merck, UK) from *Saccharomyces cerevisiae*, as previously described ^9^. Mice received an IP injection of the vehicle (PBS; Sigma-Aldrich), or PEPITEM, its tripeptide pharmacophores or their peptidomimetics, concurrently with zymosan administration, and again at the 24-h time-point. Researchers were blinded to treatment groups and animals were allocated randomly. After 48 h, mice were sacrificed by terminal anaesthesia (isoflurane) and conformation by cervical dislocation and tissues were collected as described below.

### 2.9 Flow cytometry on peritoneal lavage fluid

The peritoneal cavity was lavaged with ice-cold 5 mM EDTA. Peritoneal lavage fluid (PLF) was centrifuged at 400 *g* for 5 min: supernatant was stored at −80 °C and cells were resuspended in MACS buffer (0.1 mM EDTA, 0.6% BSA in PBS, all from Sigma-Aldrich). Samples were blocked with FcR blocker (Miltenyi Biotec) prior to staining with the following antibodies for 20 min at 4 °C, after which samples were washed and fixed with 2% formaldehyde: anti-CD3 PECy7 (clone 145-2c11), anti-CD4 eFluor450 (clone GK1.5), anti-CD8 PE-TexasRed (clone 5H10), anti-CD45 APC-CY7 (clone 104), F4/80 FITC (clone BM8), Ly6G APC (clone 1A8; all from BD). Compensation controls were generated using cells isolated from the spleen. Immediately prior to analysis CountBright beads (Invitrogen) and Zombie Aqua (Biolegend) were added and samples were acquired using Fortessa-X20. Data were analysed offline using FlowJo (V-10.2.6) using the flow cytometry gating strategy previously described ^9^.

### 2.10 Imiquimod-induced psoriasis mouse model

Two days before the initiation of the study, mice were shaved along their backs with electric clippers, and a depilation lotion (Nair; Church & Dwight Co., Ewing, NJ) was applied to completely remove hairs ^23^. For the establishment of psoriasis mouse model, mice were topically treated with 62.5 mg of 5% imiquimod (IMQ) cream (corresponding to 3.125 mg of the active compound) on shaved skin daily for 7 consecutive days. For control (CTRL) animals, vehicle cream (Vaseline, Sigma-Aldrich; 16415) was used in the same amount ^24, 25^. PEPITEM and SVT[NH-Ethyl] (100 μg mL^−1^) were administrated topically concurrently, applying the imiquimod cream over 7 days, ensuring the same areas were treated daily. The severity of back skin was assessed by erythema, scaling and thickness according to the clinical Psoriasis Area and Severity Index (PASI) as previously described ^24^. Briefly, erythema, scaling, and thickening were daily scored independently by three investigators who were blinded to the experimental conditions, on a scale from 0 to 4: 0, none; 1, slight; 2, moderate; 3, marked; 4, very marked ^26^. The cumulative score (erythema plus scaling plus thickening) served as a measure of the severity of inflammation ^27^. Skin, skin-draining lymph node and spleen samples were harvested at the experimental endpoint (day 7), and length was measured and weighed. The obtained samples were stored at −80 °C and used for subsequent experiments ^28, 29^.

### 2.11 Histology

For haematoxylin and eosin (H&E; Carlo Erba, Italy) staining, skin samples were excised immediately, thoroughly rinsed with physiological saline solution and fixed in 4% formaldehyde prior to being embedded in paraffin ^24, 29^. Sections of 5-6 μm were cut and deparaffinized with xylene before the staining was performed ^15, 21^. In all cases, a minimum of ≥ 3 sections per animal were evaluated. Images were taken by a Leica DFC320 video camera (Leica, Milan, Italy) connected to a Leica DM RB microscope using the Leica Application Suite software V2.4.0. and analysed using the Image Pro image analysis software package.

### 2.12 Isolation of immune-infiltrated cells

Spleens were mechanically dissociated to obtain single-cell suspensions, as described ^22, 30^. For single-cell suspensions from skin, 2×3 cm sections were washed two times with Hanks′ balanced salt solution (HBSS) to remove the remaining IMQ cream, minced with sharp scissors, and incubated with Dispase II solution (5 mg mL ^−1^ in PBS, Sigma-Aldrich; D4693) in a 12-well plate at 37 °C for 1 h. Subsequently, skin samples were transferred in a 6-well plate and digested with Dermis Dissociation Buffer made with DMEM, 100 mg mL^−1^ DNase I (Sigma-Aldrich; DN25) and 1 mg mL^−1^ Collagenase (Sigma-Aldrich; C2139), at 37°C for 1 h with continuous stirring. Then, digested skin samples were incubated for an additional 30 min in a cell culture incubator and, after repeated pipetting, filtered through 40-μm cell strainer ^31^. All cell suspensions were treated with red blood cell lysis buffer and washed in PBS for total cell count before flow cytometry analysis. Cell number was determined by TC20 automated cell counter (Bio-Rad, Milan, Italy) using Bio-Rad’s disposable slides and TC20 trypan blue dye (Product Number:1450013; Bio-Rad; 0.4% trypan blue dye w/v in 0.81% sodium chloride and 0.06% PBS) ^15, 20^.

### 2.13 Flow cytometry analysis

Cells collected from digested psoriatic skin tissues and splenocytes were washed in buffer (PBS containing 1% BSA and 0.02% NaN_2_) and incubated with eBioscience^™^ Fixable Viability Dye eFluor 780 (Thermo Fisher Scientific; 65-0865-14) before the staining with fluorescence-conjugated antibodies. Specifically, skin-digested cells were stained with anti-LY6G APC (clone 1A8; BioLegend) and anti-LY6B.2 [7/4] FITC (clone 7/4; Abcam) for 60 min at 4 °C to define neutrophils and monocyte as LY6B.2(7/4)^+^LY6G^high^ and LY6B.2(7/4)^+^LY6G^low^ respectively ^32^. Splenocytes were surface stained with the following conjugated antibodies (all from BioLegend, London, UK): anti-CD4 APC (clone GK1.5), anti-CD3 PE (clone 17A2) and anti-CD8 FITC (clone 53-6.7) for 60 min at 4 °C. After washing, cells were fixed, permeabilized, and stained intracellularly with anti-IFN-γ PE (clone XMG1.2) and anti-IL-17A PE antibody (clone TC11–18H10.1). T-helper (Th)1 and Th17 populations were defined as CD4^+^IFN-γ^+^ and CD4^+^IL-17^+^ cells respectively, according to the flow cytometry procedure previously described ^15, 22^. At least 1 × 10^4^ cells were analysed per sample, dead cells were excluded using cell viability marker, and positive and negative populations were determined based on the staining obtained with related IgG isotypes. Flow cytometry was performed on BriCyte E6 flow cytometer (Mindray Bio-Medical Electronics, Nanshan, China) using FlowJo software operation ^20, 33^. The gating strategy is described in **Supplementary Fig. 2A-C**.

### 2.14 Cytokine and chemokine protein array

Cells isolated from digested skin tissue were lysed with lysis buffer supplemented with Complete™ Protease Inhibitor Cocktail (Roche; 04 693 116 001) ^15^. After centrifugation (4500 *g* at 4 °C) for 10 min supernatants were collected, quantified using a Bradford protein method and assayed for proteome profiler cytokine array kit (R&D Systems; ARY006) ^15^. According to the manufacturer’s instructions, equal volumes (1.5 mL) of the pulled skin-cells lysates from all experimental conditions were then incubated with the pre-coated proteome profiler array membranes. Dot plots were detected by using the enhanced chemiluminescence detection kit and Image Quant 400 GE Healthcare software (GE Healthcare, Italy) and successively quantified using GS 800 imaging densitometer software (Biorad, Italy) ^15, 34^.

### 2.15 Multiplex cytokine assay

For isolation of lymphocytes, skin-draining lymph nodes were cut into small pieces, mechanically dissociated and filtered through 70-mm nylon cell strainer ^30^. Obtained single-cell suspensions were lysed of red blood cells and then purified using Ficoll-Paque density gradient centrifugation (1025 *g* at 20 °C) for 20 min with slow acceleration and brakes turned off. The formed cell layer was washed in PBS (300 *g* for 10 min at 20 °C) and resuspended in RMPI/10% FBS/1% penicillin-streptomycin. Purified lymphocytes (1 x 10^6^ mL^−1^ per well) were then seeded in 24-well plates and stimulated with phorbol myristate acetate (PMA; 0.5 μg mL^−1^, Sigma-Aldrich.) and Ionomycin (IONO; 0.25 μg mL^−1^, Sigma-Aldrich) ^35^. After 24 h, supernatants were collected, and the concentrations of Th-related cytokines were determined using the multi-LEGENDplex™ analyte flow assay kit (mouse Th Panel (12-plex); Biolegend; 741043). Briefly, antibodies specific to the 12 analytes were conjugated to 12 different fluorescence-encoded beads. The beads were mixed with the supernatants, incubated for 2 h at room temperature (RT), washed, and incubated for 1 h with detection antibodies. Finally, streptavidin-PE was added and incubated for 30 min, and the beads were washed and acquired using BriCyte E6 flow cytometer (Mindray Bio-Medical Electronics). The results were analysed by using the Legendplex software (version 8.0) ^36^. The gating strategy is shown in **Supplementary Fig. 3**.

### 2.16 Western blot analysis

Skin lesions homogenates were subjected to SDS-PAGE (10% gel) using standard protocols ^37, 38^. The proteins were transferred to nitrocellulose membrane (0.2-μm nitrocellulose membrane, Trans-Blot^®^TurboTM, Transfer Pack, Bio-Rad Laboratories, Hercules, CA, USA, RRID: SCR_008426) in transfer buffer (25-mM Tris–HCl pH 7.4 containing 192-mM glycine and 20% v/v methanol) at 400 mA for 2 h. Membranes were incubated for 2 h with non-fat dry milk (5% w/v) or BSA (4.5% w/v) in PBS supplemented with 0.1% (v/v) Tween 20 (PBS-T) at room temperature and then incubated at 4 °C with the appropriately diluted primary antibodies overnight: rabbit monoclonal NF-kB p65 (1:1000; D14E12), rabbit monoclonal COX-2 (1:1000; D5H5; both from Cell Signaling) and mouse monoclonal anti-beta-actin (0.01 µg mL^−1^; MAB8929; R&D Systems). After lavages with PBS-T, blots were incubated with a 1:3000 dilution of related horseradish peroxidase-conjugated secondary antibody (Donkey anti-Mouse IgG Heavy and Light Chain Antibody HRP Conjugated, A90-137P or Sheep anti-Rabbit IgG Heavy and Light Chain Antibody HRP Conjugated, A120-100P; Bethyl Laboratories) for 2 h at RT and finally washed 3 times with PBS-T. Protein bands were detected by using the enhanced chemiluminescence method (Clarity™ Western ECL Substrate, BioRad Laboratories, USA) and Image Quant 400 GE Healthcare software (GE Healthcare, Italy). Bands were quantified using the GS 800 imaging densitometer software (Biorad, Italy) and normalized with respective beta-actin ^22, 39, 40^. Uncropped original western blots are presented in **Supplementary Fig. 4A-C**.

### 2.17 Data and statistical analysis

Statistical analysis complies with the international recommendations on experimental design and analysis in pharmacology and data sharing and presentation in preclinical pharmacology ^15, 40^. Data are presented as means ± S.D. Normality was tested prior to analysis with one or two-way ANOVA followed by Bonferroni’s or Dunnett’s for multiple comparisons, where p ≤0.05 was deemed significant. GraphPad Prism 8.0 software (San Diego, CA, USA) was used for analysis. Animal weight was used for randomisation and group allocation to reduce unwanted sources of variations by data normalisation. No animals and related *ex vivo* samples were excluded from the analysis. *In vivo* studies were carried out to generate groups of equal size (n=6 of independent values), using randomisation and blinded analysis. Data was considered significant at p<0.05,

## 3 Results

### 3.1 Identifying functionally active pharmacophores within the PEPITEM sequence

In the early phase of our drug a discovery programme, we synthesised 13 novel peptides with alanine substitutions progressively along the length of the 14 AA PEPITEM sequence SVTEQGAELSNEER (**Supplemental Table 1**). The synthesis of peptides and their mimetics for optimising ADME characteristics is expensive and time consuming and working with the smallest sequence that retains all relevant biological functions is desirable. An alanine sweep can be used to identify such motifs. In this strategy each amino acid in the sequence is substituted sequentially with alanine, and the resulting peptides screened for activity in a relevant assay to identify those that are functionally redundant. As AA7 is an alanine in the native PEPITEM sequence, we used this is a positive control for regulation of lymphocyte transmigration across TNF-α/IFN-γ-stimulated endothelial cells. Surprisingly all the peptides inhibited lymphocyte migration across cytokine stimulated endothelial cells to a similar degree, when compared to native PEPITEM (**Fig. 1A**).

**Figure 1.**
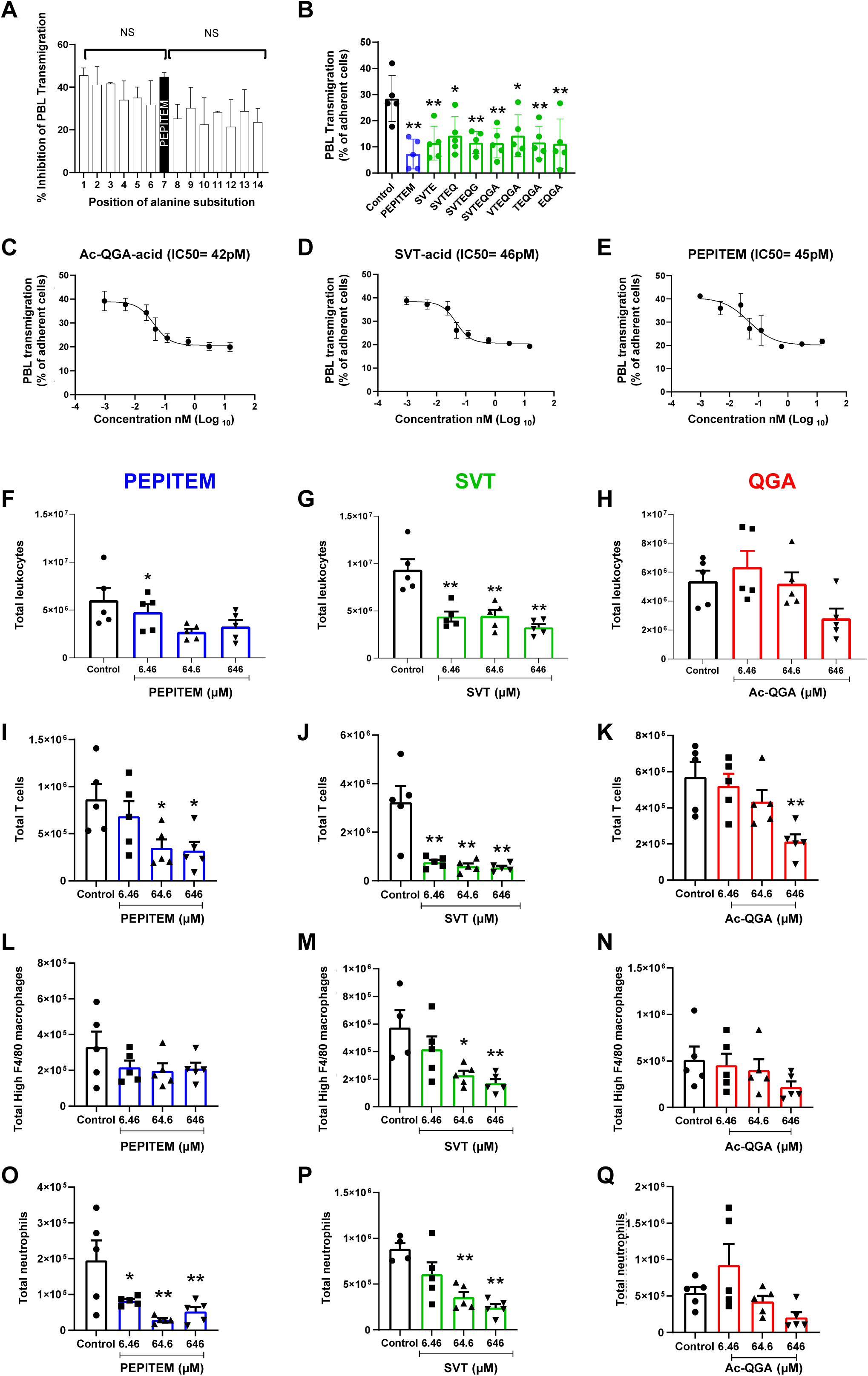
Identification of the functional motifs within the PEPITEM sequence and their efficacy *in vitro* and *in vivo*. **(A-E)** PBL were pre-treated with peptides (20ng/ml) for 5 minutes, prior to addition to cytokine-stimulated endothelium. PBL transmigration was assessed at 20 minutes and expressed as (A) % inhibition or (B) % of adherent cells. **A)** Effects of an alanine sweep on the function of PEPITEM analysed by one way ANOVA with Dunnett’s test with comparisons made to the effects of PEPITEM. **B)** The effects of different truncated peptides from the SVTEQGA sequence. Data are mean ± SEM, n=3-6. NS=not statistically significant; * = p< 0.05, ** = p< 0.01 analysed by one way ANOVA with Dunnett’s test with comparisons made to untreated control. Dose-response curves for **C)** QGA, **D)** SVT and **E)** PEPITEM (F-Q) Mice were treated with three doses of PEPITEM, SVT or QGA at the same point as induction of peritonitis by zymosan. **F), G) and H)** the effects of peptides on total peritoneal leukocyte counts; **I,), J) and K)** the effects of peptides on total peritoneal T cell counts. **L), M) and N)** the effects of peptides on total peritoneal macrophage counts. **O), P) and Q)** the effects of peptides on total peritoneal neutrophil counts Data are mean ± SEM from n=5 animals in each group. * = p< 0.05, ** = p< 0.01 analysed by one-way ANOVA with Bonferroni’s posthoc-test.

A logical hypothesis for these observations is an inbuilt functional redundancy within PEPITEM, in other words more than one active peptide sequence within PEPITEM was operating to regulate T-cell trafficking. Cutting the molecule in half delivered a heptapeptide sequence (SVTEQGA) with full activity, i.e. could significantly reduce lymphocyte trafficking *in vitro* (**Fig. 1B**). This demonstrated that the active sequences resided within the heptapeptide SVTEQGA. Thus, we truncated SVTEQGA from either end by sequentially removing amino acids. This revealed the minimum active sequences within the heptapeptide capable of significantly reducing lymphocyte migration were the tripeptides SVT and QGA (synthesised as SVT-acid and ac-QGA-acid for technical reasons but referred to as SVT and QGA going forward) (**Fig. 1B**). Despite dissimilar structures (**Supplementary Table 2**), SVT and QGA significantly reduced lymphocyte migration *in vitro* to a similar degree as PEPITEM, with equivalent IC50’s in the range of 40-50 pM (**Fig. 1C-E**). To validate the *in vitro* dose response data, we utilised zymosan induced peritonitis to model acute inflammation. The trafficking dynamics of various leukocyte populations was established at 48 h after administration of zymosan using flow cytometry of labelled leukocytes collected by peritoneal lavage. We found PEPITEM reduced total leukocyte counts (CD45^+^) at the two highest doses, and significantly at 64.6 nM (**Fig. 1F**). Strikingly, SVT caused a significant reduction of CD45^+^ leukocytes across all doses, whilst QGA only showed effects at the highest dose (**Fig. 1G and H**). Consistently with previous publications ^6, 7, 8, 9^ all three peptides reduced CD3^+^ T cells at the highest dose, with SVT and PEPITEM also showing significant inhibitory effects at lower dose (**Fig. 1I-K**). Whilst, SVT significantly reduced F4/80^high^ macrophages at all concentrations, PEPITEM and Ac-QGA had no effect (**Fig. 1L-N**). A dose dependent reduction in peritoneal Ly6G^+^ neutrophil numbers was also observed when mice were treated with PEPITEM and SVT, and to a lesser extent with QGA (**Fig. 1 O-Q**). Taken together, the tripeptides SVT and QGA show biological activity in the inhibition of leukocyte trafficking comparable to the full-length native PEPITEM.

### 3.2 Engineering peptidomimetics from SVT and QGA that have anti-inflammatory efficacy

A major drawback of utilising peptides therapeutically is their susceptibility to rapid degradation in the plasma by endogenous peptidases. This can be readily assessed by adding peptides to human or murine plasma and assaying the rate of decline of the native sequence by mass spectrometry. We found that PEPITEM was remarkably stable showing no discernible enzymatic degradation over 180 min, however, the tripeptides were less stable (**Supplementary Table 2**). In addition, molecules of less than 50kDa in MW are cleared rapidly from the circulation by renal filtration (PEPITEM has a MW of approx. 1.5kDa). For example, upon intravenous (IV) injection of a tritiated version of PEPITEM (SVTEQGAE-[4,5-^3^H-Leu]-SNEER-acid) and scintillation counting of collected blood, we observed a rapid loss (within 2 min) to approximately 5% of the original signal (**Supplementary Fig. 5A**). PEPITEM showed longer residence times in other organs when a fluorescently labelled version (Alexa Fluor-680; AF680) was used to track peptide residence in different tissues after IV injection (**Supplementary Fig. 6B**). There were small but significant depots of labelled peptide resident in the heart, lungs and spleen. However, by far the largest signal was found in the kidneys implying a preferential assimilation by these organs for elimination in the urine (**Supplementary Fig. 5B and C**).

As the tripeptides were functional both *in vitro* and *in vivo* but had inferior physical characteristics to PEPITEM, we began a programme of iterative peptide optimisation using N and C terminal modifications and substitution of AA with none-natural variants, including d-isomeric forms, which are not recognised by degradative enzymes. The process followed the workflow detailed in **Supplementary Fig. 6**, where peptides were designed, synthesised by standard solid state peptide synthesis methodology, and then assessed for bioactivity using the in *vitro* lymphocyte migration assay. The 8 peptides showing efficacy equivalent or greater than PEPITEM were tested at multiple concentrations to determine IC50 (**Supplementary Table 2**). Simultaneously, 17 compounds were tested for stability in plasma to establish the effects of specific structural changes on this parameter (**Supplementary Table 2**). The calculated physicochemical properties of the peptides and peptidomimetics (using Optibrium StarDrop version 7.6.1 software) are listed in **Supplementary Table 3**. Based on efficacy *in vitro* and stability in plasma, two peptidomimetics were taken forward for further study: [pGlu]-GA-amide and SVT[NH-Ethyl] which have modifications incorporated to improve metabolic stability to proteases. The selected peptidomimetics showed approximately equal efficacy *in vitro* to the native peptides, significantly reducing migration of lymphocytes across inflamed endothelial cells (**Fig. 2A**). Similarly, both mimetics significantly reduced total CD45^+^ leukocyte levels, including total CD3^+^T cells, F4/80^high^ macrophages and Ly6G^+^ neutrophils numbers within the inflamed peritoneum *in vivo* (**Fig. 2B-I**). Thus, peptidomimetics were capable of regulating leukocyte trafficking with the at least the same efficacy as PEPITEM and the native tripeptides.

**Figure 2.**
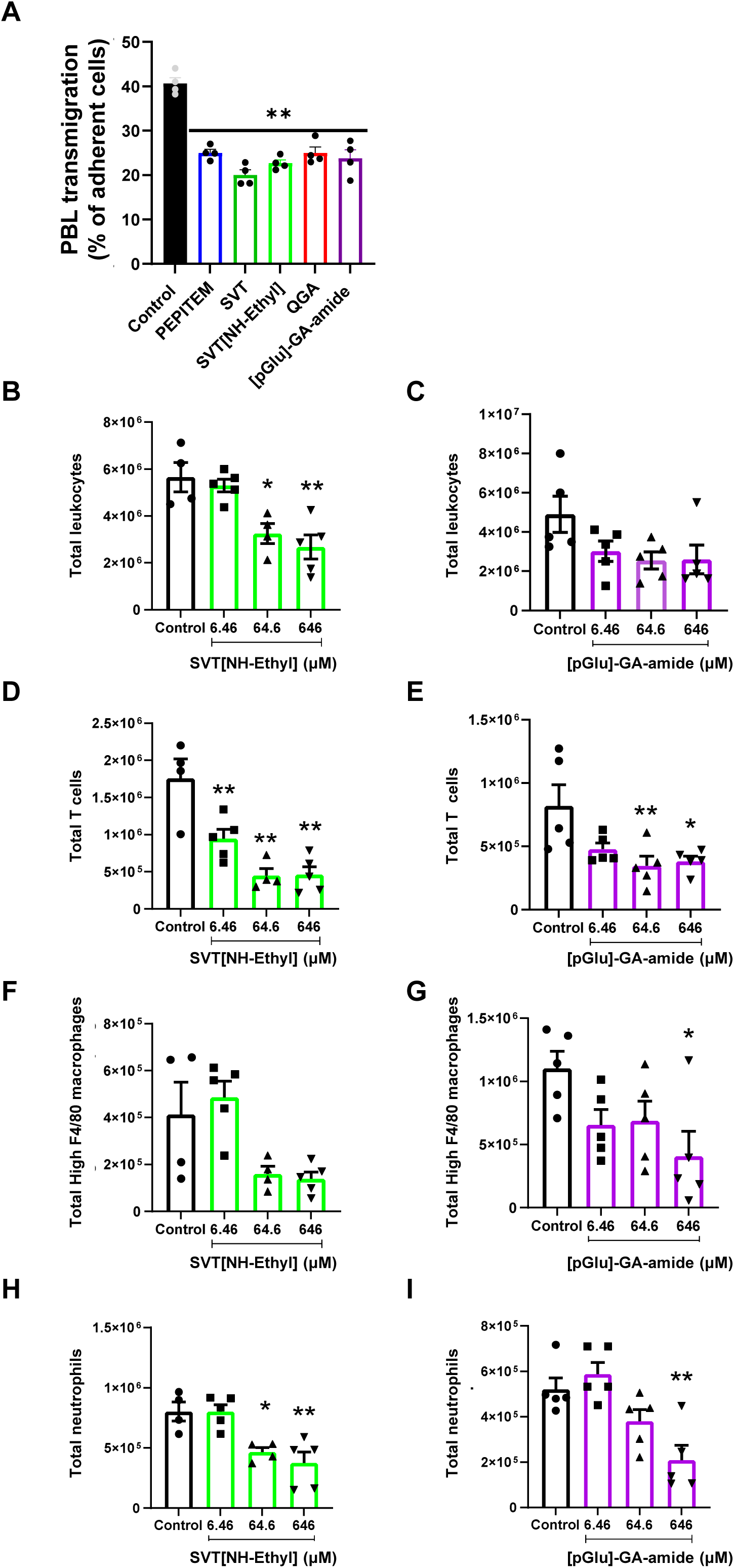
The effects of peptidomimetic sequences on leukocyte trafficking *in vitro* and *in vivo*. **A)** The effects of PEPITEM, SVT, QGA, SVT[NH-Ethyl] and [pGlu]-GA-amide on lymphocyte trafficking *in vitro* where PBL were pre-treated with peptides (20ng/ml) for 5 minutes prior to addition to cytokine-stimulated endothelium. PBL transmigration was assessed at 20 minutes. Data is mean ± SEM of 4 independent experiments; ** = p< 0.01 analysed by one-way ANOVA with Bonferroni’s posthoc-test. The dose-dependent effects of SVT[NH-Ethyl] and [p-GLU]-GA-amide on leukocyte populations in the peritoneum of zymosan treated mice. **B) and C)** the dose dependent effects on total peritoneal leukocytes **D) and E)** The dose-dependent on total peritoneal CD3^+^ T **F) and G)** The dose-dependent effects on total peritoneal F4/80^high^ macrophages **H) and I)** The dose-dependent effects on total peritoneal neutrophils, in a model of zymosan-induced peritonitis. **B-I**, data are mean ± SEM from n=5 animals in each group. * = p< 0.05, ** = p< 0.01 analysed by one-way ANOVA with Bonferroni’s posthoc-test.

### 3.3 Topical delivery of peptides can reduce local inflammation and disease severity in a model of plaque psoriasis

PEPITEM improves clinical severity and reduces leukocyte trafficking in several models of immune-mediated inflammatory disease when administered systemically ^6, 7, 8, 9^. To further investigate the therapeutic efficacy of the mimetics *in vivo* we trialled peptides by topical application in a preclinical model of plaque psoriasis. This mode of delivery is so far untested for agents of the PEPITEM pathway. Plaque psoriasis was induced by topical application of imiquimod in an emollient cream ^24, 25^ that induced psoriasis within 48 h, with disease becoming progressively worse over 7 days (**Fig. 3A and B**). We selected a single lead agent based on activity *in vitro* and *in vivo* (i.e. SVT[NH-ethyl]) that was tested therapeutically by mixing into the imiquimod cream for simultaneous topical application every day for 7 days. Visual and histological assessment of skin showed a clear reduction in disease in the presence of PEPITEM and SVT[NH-ethyl] when compared to untreated psoriasis (**Fig. 3A and B**). Disease was formally assessed using PASI score (**Fig. 3C**), where erythema (redness; **Fig. 3D**), desquamation (scaling; **Fig. 3E**) and induration (thickening; **Fig. 3F**) were each assessed on a scale of 0-4, to give a combined maximal score of 12. PEPITEM and SVT[NH-ethyl] significantly reduced the PASI score by ~50% when compared to untreated psoriasis group (which is a clinically significant intervention). To assess the effect of the peptides on immune infiltration at day 7, skin was digested and analysed by flow cytometry. As expected, disease onset significantly increased neutrophil and monocyte infiltration into the skin compared to untreated control animals (**Fig. 3G and H**). Treatment with either PEPITEM or SVT[NH-ethyl] significantly reduced monocyte and neutrophil levels, where SVT[NH-ethyl] was significantly superior to PEPITEM in regulating the neutrophilic infiltrate (**Fig. 3G**). Gating strategy is reported in **Supplementary Fig. 2A.** Taken together, this data demonstrates that local topical application of the mimetics reduces disease severity through regulating leukocyte infiltration.

**Figure 3.**
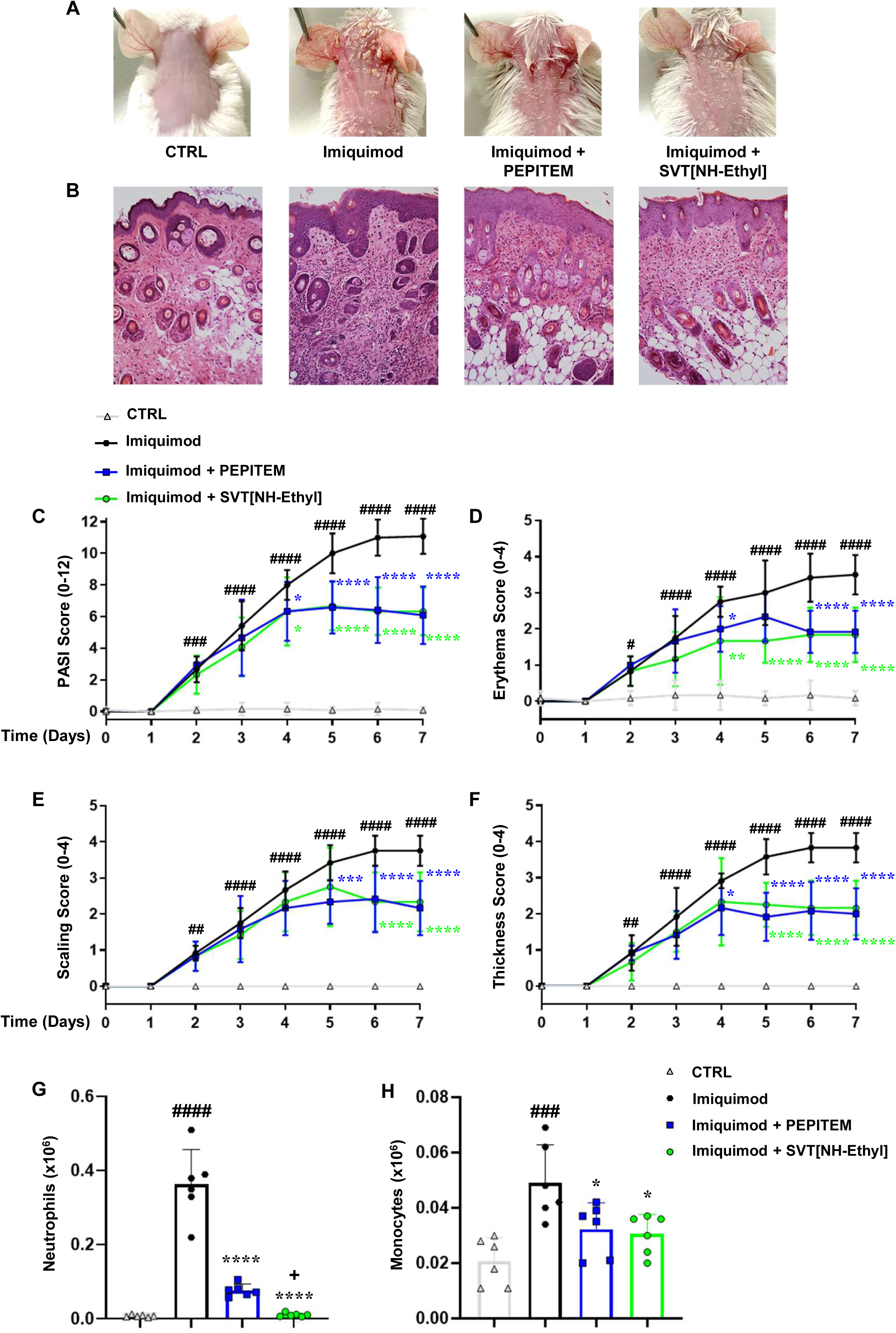
PEPITEM and SVT[NH-ethyl] relieve local inflammation and disease severity in a model of Imiquimod (IMQ)-induced psoriasis. **A)** Representative photographs of skin damage from mice treated with IMQ or vaseline (CTRL) in presence or absence of PEPITEM and SVT[NH-Ethyl] (both administered at 100 μg ml^−1^ daily, with topical application). **B)** H&E staining of skin sections from the different experimental conditions at the experimental endpoint (day 7). **C)** Scoring time-line curves of back skin cumulative PASI score, determined by calculating single **D)** erythema, **E)** scaling and **F)** thickness over time for 7 days. Flow cytometry analysis of digested skin tissues applied to determine **G)** neutrophils (LY6B.2(7/4)^+^LY6G^high^) and **H)** monocytes (LY6B.2(7/4)^+^LY6G^low^), pre-gated on living cells (see flow cytometry strategy reported in Supplementary Fig. 2A). Histograms indicate the total positive populations (expressed as x10^6^) in the different experimental conditions. Data are presented as means ± S.D. of n=6 animals in each group. Statistical analysis was conducted by one- or two-way ANOVA followed by Bonferroni’s or Dunnett’s posthoc-test. ^#^P ≤ 0.05, ^##^P ≤ 0.01, ^###^P ≤ 0.001, ^####^P ≤ 0.0001 *vs* CTRL group; *P ≤ 0.05, **P ≤ 0.01, ***P ≤ 0.001, ****P ≤ 0.0001 *vs* Imiquimod group; ^+^P ≤0.05 *vs* PEPITEM group.

### 3.4 Topical application of peptides regulates the local dermal microenvironment

To further investigate the molecular mechanisms underpinning bioactivity of the mimetics, we explored changes in the composition of the inflammatory mediators within the skin that are responsible for driving leukocyte infiltration using a proteome profiler cytokine array (**Fig. 4A**). Of the 32 analytes measured, 22 were induced *de novo*, and a further 8 were upregulated from basal levels in response to psoriasis (**Fig. 4A and B**). The impact of the mimetics on these analytes was complex: following PEPITEM or SVT[NH-Ethyl] treatment, expression of 8 analytes (BLC, IL-3, IL-4, IL-23, IP-10, I-TAC, MIP-2 and TARC) were no longer detectable and a further 7 analytes (C5a, IL-1α, IL-17, INF-γ, IL-27, IL-1ra and TREM-1) were reduced (**Fig. 4A and B**). Interestingly, PEPITEM treatment reduced expression of 3 analytes (IL-13, MIG and TIMP-1) that remained unaffected by SVT[NH-Ethyl], whilst SVT[NH-ethyl] diminished expression of 9 analytes (G-CSF, GM-CSF, M-CSF, Eotaxin, KC, MCP-5, MIP-1β, RANTES and TNF-α) that were unaffected by PEPITEM treatment (**Fig. 4C**). The wide-ranging reduction in chemokines, cytokines and growth factors released into the inflamed skin implies that PEPITEM and its mimetics have powerful regulatory control of the synthesis and secretion of these agents, and thus can dramatically influence inflammatory processes.

**Figure 4.**
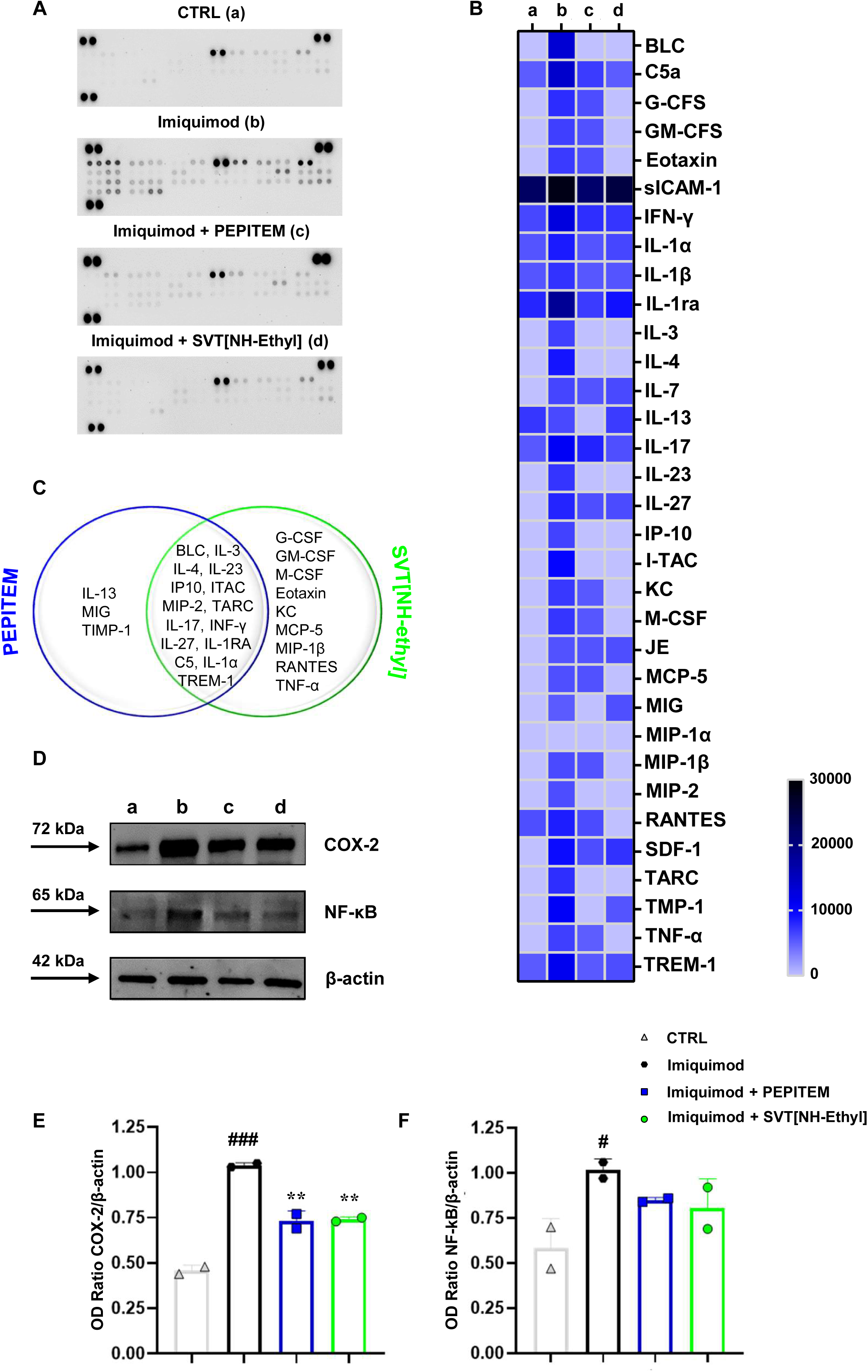
PEPITEM and SVT[NH-Ethyl] regulate cyto-chemokine levels in inflamed skin through the regulation of COX-2/NF-κB signaling. **A)** Proteome profiler cytokine array for CTRL, Imiquimod (IMQ) alone or in combination with PEPITEM or SVT[NH-Ethyl] groups performed on digested skin lysates. **B)** Densitometric analysis shown as a heatmap and expressed as INT/mm^2^. Data are presented as means ± S.D. (colormap: double gradient) of n=6 animals in each group. **C)** Stacked Venn diagrams of cyto-chemokines up-regulated by IMQ and significantly modulated by the treatment with PEPITEM and SVT[NH-Ethyl] (both administered at 100 μg mL^−1^ daily, with topical application). **D)** Western blot for COX-2, NF-κB and β-actin performed on skin lesions homogenates from different experimental conditions as technical duplicate of one experiment run with n=6 mice per group pooled. **E) and F)** Cumulative densitometric values are expressed as OD ratio with β-actin and presented as means ± S.D. of n=6 animals per group pooled. Statistical analysis was conducted by one- or two-way ANOVA followed by Bonferroni’s or Dunnett’s posthoc-test. ^#^P ≤ 0.05, ^###^P ≤ 0.001 *vs* CTRL group; **P ≤ 0.01 *vs* Imiquimod group.

Next, we investigated the expression of the key up stream mediators responsible for regulating cytokine production, nuclear factor kappa B (NF-kB) and cyclooxygenase-2 (COX-2). Upon induction of psoriasis both NF-kB-p65 and COX-2 were upregulated significantly compared to untreated control animals, when assessed by western blotting (**Fig. 4D**). PEPITEM and SVT[NH-ethyl] significantly reduced the expression of COX-2, and both peptides tended to reduce NF-kB-p65, albeit this remained non-significant (**Fig. 4D-F**) (uncropped original western blots are reported in **Supplementary Fig. 4A-C**). Thus, PEPITEM and derived peptides regulate chemokine and cytokine expression in inflamed skin most likely through regulation of mediators known to support their induction, transcription and secretion.

### 3.5 Topical application of PEPITEM and its pharmacophores alter systemic inflammatory responses in a model plaque psoriasis

Previous characterisation of the IMQ-induced psoriasis model showed that topical application to the skin causes both local (skin) and systemic changes in inflammatory leukocyte populations in lymphoid organs ^41, 42, 43^. Notably here we observed a significant splenomegaly (**Supplementary Fig. 7**), accompanied by an increase in Th1 and Th17 cells (gating strategy is reported in **Supplementary Fig. 2B and C**). Spleen length, weight and weight/body weight ratio were significantly increased upon induction of disease (**Fig. 5A** and **Supplementary Fig. 7**). Administration of PEPITEM and SVT[NH-Ethyl] significantly reduced all these parameters (**Fig. 5A and Supplementary Fig. 7**). Psoriasis induced splenomegaly was associated with an expansion of both CD4^+^ and CD8^+^ lymphocyte pools (**Fig. 5B and C**), as well as a marked increase in Th1 (IFN-γ^+^) and Th17 (IL-17^+^) cells (**Fig. 5D and E**), which were significantly reduced upon treatment with PEPITEM and SVT-[NH-ethyl] (**Fig. 5B-E**). The systemic immunological imbalance was also evaluated examining the functional responses of T-cells isolated from skin draining lymph nodes (following *ex vivo* stimulation with PMA and IONO) (gating strategy is reported in **Supplementary Figure 3**). We observed induction of effector T-cell cytokines by psoriasis, specifically elevation of 7 analytes (IFN-γ, TNF-α, IL-2, IL-6, IL-10, IL-17A and IL-22) (**Fig. 5F**). Co-administration of imiquimod with PEPITEM or SVT[NH-ethyl] significantly reduced the expression of IL-6, TNF-α and IL-17A (**Fig. 5G-I**). Collectively these data demonstrate that topical application of PEPITEM and its mimetics on psoriatic skin not only moderates inflammation and disease locally but had significant effects on systemic inflammatory and immunological responses induced by disease.

**Figure 5.**
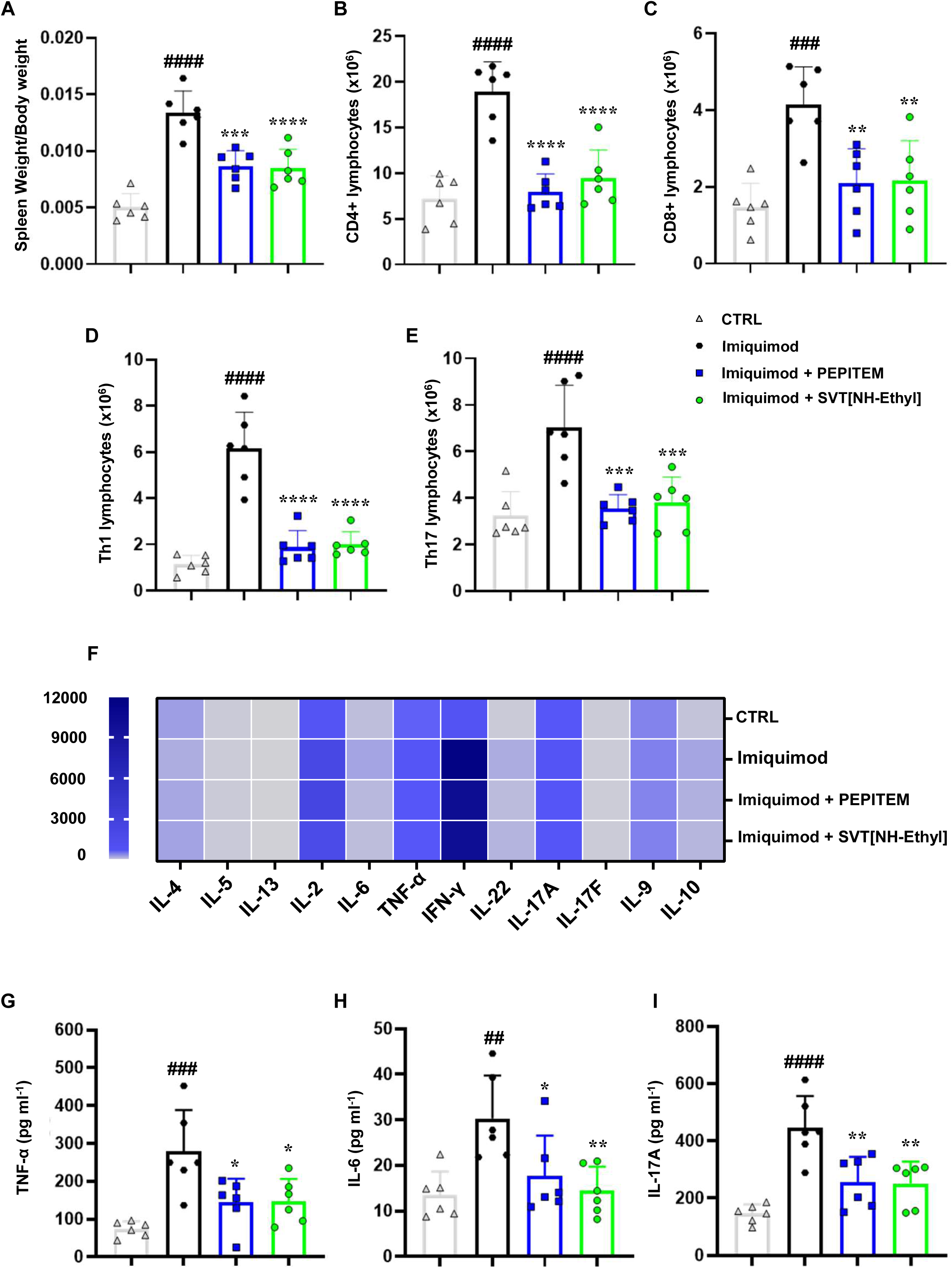
The effect of PEPITEM and SVT[NH-Ethyl] is mirrored in the lymphoid organ repertoire. **A)** Spleen weight/mouse body weight ratio (expressed as g) evaluated at the experimental endpoint (day 7). Flow cytometry analysis of splenic **B)** CD4^+^, **C)** CD8^+^, **D)** Th1 (CD4^+^IFN-γ^+^) and **E)** Th17 (CD4^+^IL-17^+^) subsets, pre-gated on living cells (see flow cytometry strategy reported in Supplementary Fig. 2B and C). Histograms indicate the total positive populations (expressed as x10^6^) in the different experimental conditions. **F)** Quantification of T-related cytokines performed by a multi-LEGENDplex^™^ analyte flow array (gating strategy is reported in Supplementary Fig. 3) on lymphocytes isolated from skin-draining lymph nodes, following *ex vivo* stimulation with PMA and ionomycin, presented as a heatmap (colormap:double gradient) and expressed as pg mL^−1^. **G), H) and I)** TNF-α, IL-6 and IL-17A cytokines extrapolated from the heatmap and represented graphically as histograms. Data are presented as mean ± S.D. of n=6 animals in each group. Statistical analysis was conducted by one- or two-way ANOVA followed by Bonferroni’s or Dunnett’s posthoc-test. ^##^P ≤ 0.01, ^###^P ≤ 0.001, ^####^P ≤ 0.0001 *vs* CTRL group; *P ≤ 0.05, **P ≤ 0.01, ***P ≤ 0.001, ****P ≤ 0.0001 *vs* Imiquimod group.

### 3.6 *In vitro* mechanistic studies on the actions of PEPITEM, tripeptides and peptidomimetics on cells of the skin stromal environment

Psoriasis is driven by multiple cell types within the skin, with fibrosis/hyperplasia driven by the fibroblasts, keratinocytes, and tissue-resident macrophages^13^. Moreover, tissue-resident stromal cells act as a major source of inflammatory mediators, driving leukocyte infiltration ^13^. Given the modulation of the dermal microenvironment by the peptides and peptidomimetics, we sought to investigate the impact of these agents on modulating soluble mediator release with *in vitro* cellular models of the skin stromal environment using macrophage, fibroblast and keratinocyte cell lines. Moreover, the higher throughput nature of the assay allowed us to compare a greater range of peptides. Using an MTT assay we showed a lack of cytotoxicity of these peptides up to 30 ng mL^−1^ at both 4 and 24 h (**Fig. 6A and B**). We then showed that that LPS induced significant release of both TNF-α and IL-6 in J774A.1 macrophages and NIH-3T3 fibroblast cell lines, which was significantly inhibited by addition of PEPITEM or SVT[-NH-ethyl] but not [pGlu]-GA-amide (**Fig. 6C-F**). Of note, the native tripeptide sequences (SVT, QGA) had no significant effects on cytokine release compared to the untreated LPS. Using a human epidermal keratinocyte line (HaCaT), we tested the ability of peptides to inhibit cellular proliferation in response to the gold standard stimulation with M5 cytokine mix (IL-17A, IL-22, Oncostatin-M, IL-1α, TNF-α). M5 mix induced significant proliferation in cell cultures over 72 h, which was significantly reduced when cultures were pretreated with either PEPITEM, SVT[NH-ethyl], QGA or [pGlu]-GA-amide (**Fig. 6G**). Thus, PEPITEM and its derivatives regulates inflammation by multiple mechanisms, including effects on endothelial cells and leukocyte trafficking, as well as regulation of the induction and release of inflammatory cytokines in the stroma of the affected tissue.

**Figure 6.**
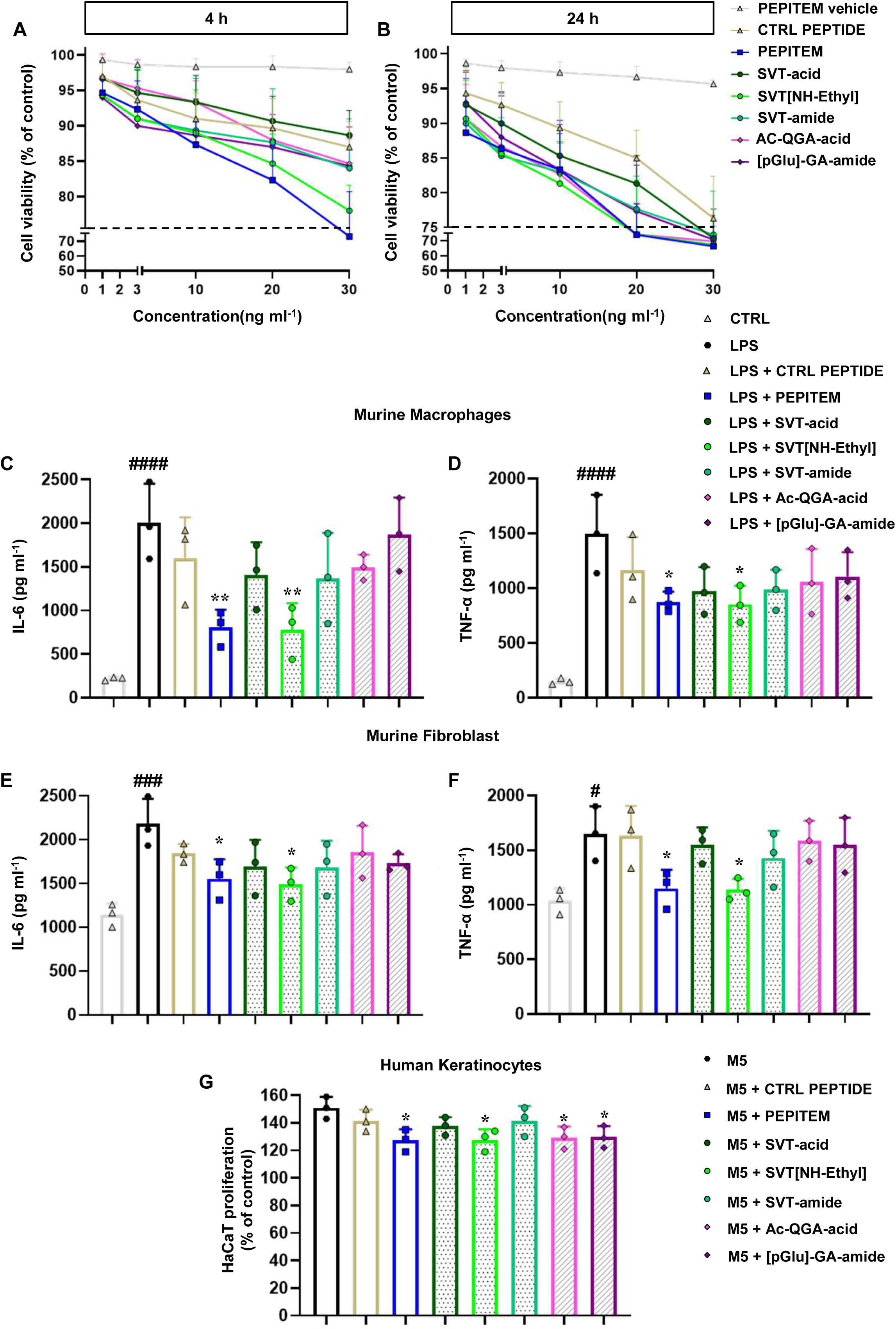
**A) and B)** *In vitro* mechanistic studies on PEPITEM, tripeptides and peptidomimetics. *In vitro* cytotoxic examination, evaluated by MTT assay, for PEPITEM, tripeptides and peptidomimetics performed on J774A.1 murine macrophage cell lines, following 4 and 24 h of treatment with the selected concentrations (1-30 ng mL^−1^). Dotted lines indicate 75% of cell viability. Data are expressed as cell viability (% of control) and presented as mean ± S.D. of 3 independent experiments. **C) and D)** IL-6 and TNF-α ELISA assays performed on supernatant of J774A.1 pre-treated with indicated compounds at the concentration of 10 ng mL^−1^ (highest non-cytotoxic concentration) and then stimulated with LPS (10 µg mL^−1^) for 24 h. **E) and F)** IL-6 and TNF-α ELISA assays performed on supernatants of NIH-3T3 mouse fibroblasts pre-treated with indicated compounds at the concentration of 10 ng mL^−1^ and then stimulated with LPS (10 µg mL^−1^) for 24 h. (**C-F**) Data are expressed as pg mL^−1^ and presented as means ± S.D. of 3 independent experiments. Statistical analysis was conducted by one-way ANOVA followed by Bonferroni’s posthoc-test. ^#^P ≤ 0.05, ^###^P ≤ 0.001, ^####^P ≤ 0.0001 *vs* CTRL group; *P ≤ 0.05, **P ≤ 0.01 *vs* LPS group. **G)** Effect of tested compounds on hyperproliferation of M5 cytokines (TNF-α, Oncostatin M, IL-1α, IL-17 and IL-22)-stimulated HaCaT cells, as *in vitro* psoriasis-like model. HaCaT hyperproliferation (expressed as cells proliferation, % of control) was measured using MTT assay and presented as mean ± S.D. of 3 independent experiments. Statistical analysis was conducted by one-way ANOVA followed by Bonferroni’s posthoc-test. *P ≤ 0.05 *vs* M5 group.

## 4. Discussion

Here we have identified smaller functional pharmacophores within the 14AA PEPITEM sequence. Two independently active tripeptides could retard leukocyte trafficking in an *in vitro* T-cell trafficking assay, as well as an *in vivo* model of acute peritonitis. A peptide optimisation programme showed that peptidomimetics could be engineered that retained the biological function of PEPITEM and the tripeptides, with some molecules having superior efficacy when compared to the parent tripeptide sequences. Importantly, PEPITEM, tripeptides and mimetics were capable of significantly reducing inflammation and disease scores in a model of plaque psoriasis when they were delivered topically to the skin in an emollient cream. Agents achieved this by regulating the synthesis and release of cytokines and chemokines from skin stromal cells (fibroblasts and macrophages), as well as limiting keratinocyte proliferation. In addition, topical application of agents also significantly reduced Th1 and Th17 cytokines and T-cell numbers in the draining lymph nodes and spleens of treated animals. Thus, PEPITEM, its pharmacophores and peptidomimetics show powerful disease moderating capabilities which are achieved by regulating a spectrum of cellular and molecular pathways in the immune system and local tissue stroma.

Previous studies have demonstrated that the PEPITEM pathway can regulate the trafficking of inflammatory CD4 and CD8 central memory T-lymphocytes using *in vitro*, *ex vivo* and *in vivo* assays ^6, 7, 8, 9^. The paradigm supporting this is predicated on the interaction of PEPITEM with a receptor, CDH15 (M-cadherin), on vascular endothelial cells, engagement of which results in localised release of sphingosine-1-phosphate (S1P). Signalling via T-cell S1PR1 and S1PR4 then down regulates the LAF-1 (CD11a/CD11b) mediated transmigration of lymphocytes across the vascular barrier and into tissue ^6^. In the current studies, control of leukocyte trafficking was also evident from within the stromal environment of the tissues, where PEPITEM could regulate the localised transcription and secretion of inflammatory cytokines. Cytokines are responsible for upregulating endothelial adhesion receptors for leukocyte capture from blood, and chemokines activate and direct migration of adherent leukocytes into inflamed tissues. Such observations extend the PEPITEM mediated regulation of leukocyte trafficking from the endothelial cells of the intravascular environment to cells of the tissue parenchyma. This has important implications for our understanding of the pharmacokinetics of these peptides. In order to achieve their regulatory effects on resident cells, peptides must achieve efficient penetration of the stromal environment. The existence of tissue depots of active peptide (as demonstrated in this study) may thus abrogate the idea that all circulating PEPITEM (and its pharmacophores) are rapidly cleared by renal excretion or peptidase degradation and indicates that peptide pharmacokinetics and pharmacodynamics may be more complex than originally anticipated.

When looking for the minimal functional sequence within PEPITEM, we were surprised to identify two independently functioning tripeptides. Analysis *in vitro* showed that these had similar IC50’s to PEPITEM when tested in a T-cell trafficking assay (although efficacy in vivo was variable). The reason for this inbuilt redundancy is not clear, but we speculate this may be an evolutionary mechanism for ensuring that point mutations in the PEPITEM sequence do not remove complete control of aspects of the inflammatory response regulated by PEPITEM. The importance of the conservation of active motifs within PEPITEM and of the PEPITEM motif within 14.3.3.ζ is exemplified by their stability across the phylogeny. Comparison on UniProt shows there is 100% concordance between rodent and human PEPITEM sequences, and even at the level of drosophila the sequence shows 85% homology with complete conservation of the SVT motif. There is also strong concordance in the sequence of 14.3.3.ζ across the phylogenetic divide, with 98.8% homology between human and mouse proteins and 78.4% homology between human and fly. Clearly evolution does not tolerate significant mutagenesis of this protein or its biologically active peptide pharmacophores.

Although SVT and QGA were equipotent *in vitro*, this was not the case *in vivo*. Generally, SVT was as effective as PEPITEM at reducing leukocyte trafficking, expression of inflammatory mediators and reducing disease scores in a preclinical model of psoriasis. QGA, on the other hand, although having a limited ability to reduce some inflammatory parameters, was certainly not as potent as an anti-inflammatory agent. Moreover, there was evidence of some differentiation of function between the two tripeptides. In an attempt to optimise the efficacy of the tripeptide pharmacophores we generated a series of peptidomimetics based on the sequence of each. The agents taken forward for further testing had superior IC50 to PEPITEM in T-cell trafficking assays *in vitro*. However, efficacy *in vivo* largely mirrored that of the parent tripeptides, with SVT derived agents proving superior in a range of *in vitro* and *in vivo* assays to those derived from QGA. Importantly, we did identify one peptidomimetic that performed consistently better than PEPITEM and the parent tripeptide, SVT[NH-Ethyl]. Thus, these experiments provide strong proof of principle that the tripeptide pharmacophores of PEPITEM and their chemically altered derivatives can be used to intervene therapeutically in inflammatory diseases and moreover, raises the possibility that anti-inflammatory agents with some degree of targeted activity could be developed from the distinct pharmacophores.

In these studies, we chose to trial PEPITEM, its pharmacophores and their peptidomimetics in a model of plaque psoriasis. This allowed us to use a mode of delivery so far untested for these agents, i.e. direct and topical application to the inflamed tissue (skin). The agents tested could penetrate the dermis and showed efficacy in the epidermal stroma, causing a clinically significant reduction in disease of ≈50% (PASI-50). In addition, topical delivery also had some effects on systemic inflammatory process, e.g. reducing disease associated splenomegaly and limiting the expansion of the CD4, CD8, Th1 and Th17 T-cells within the spleen. Moreover, the function of T-cells in skin draining lymph nodes was also altered, with reductions in the agonist induced production of effector T-cell cytokines. As these systemic aspects of the inflammatory response typify chronic inflammation in IMIDs, this aspect of their activity is likely to be advantageous therapeutically. These observations also raise the question whether other IMIDs, e.g. arthritis, could be effectively treated by application of these agents to the skin, which may be a preferable route of dosing than injection for patients. Standard of care treatments for psoriasis already utilise several topical therapies across the disease spectrum, including corticosteroids, vitamin D analogues, retinoids and calcineurin inhibitors ^44^. In the current study we show that PEPITEM has actions that are similar to steroids, e.g. reduced leukocyte trafficking and down regulation of pro-inflammatory mediators such as cytokines. This raises the possibility of utilising these peptides in combination with licensed agents to maximise the efficacy of such treatments if synergistic activity were evident. This approach could also be steroid sparing, thereby reducing the serious side effects associate with these agents or allowing lower dosing for prolonged durations without loss of efficacy.

In summary; here we show that PEPITEM, its tripeptide pharmacophores, and tripeptide peptidomimetics regulate the inflammatory response by a number of mechanisms. These include effects on cells of the stromal environment, (in addition to those already reported in bone^45^), including regulation of the NF-kB pathway that leads to significant reduction on the synthesis and release of inflammatory cytokines and chemokines into the inflammatory site, as well as inhibition of systemic inflammatory process associated with chronicity in disease. Together, regulation of these inflammatory checkpoints leads to dampening (but not ablation) of the inflammatory response, which can have benefit when applied to immune mediated inflammatory diseases such as psoriasis.

## Acknowledgments

BA was supported by Ph.D. studentship funded by BBSRC (1090384); LP was funded by a Connect Immune & Chernajovsky Foundation Grant (22937); MC was supported by a Royal Society Dorothy Hodgkin Fellowship (DH16044). SJH was supported by a Ph.D. studentship funded by Royal Society (RGF\R1\180008). ST and AM were supported by an Innovate UK (IUK) Biomedical Catalyst Grant (4870), AJI was supported by a University of Birmingham Fellowship. Support was also provided by an MRC project grant MR/T028025/1. The National Institute of Health and Care Research (NIHR) Birmingham Biomedical Research Centre (NIHR203326) and the British Heart Foundation Accelerator (BHF) (AA/18/2/34218) have supported the University of Birmingham Institute of Cardiovascular Sciences where this research was based. The opinions expressed in this paper are those of the authors and do not represent any of the listed organisations. Work was also supported by a collaborative agreement between *ImmunoPharmaLab* at the University of Naples Federico II and the University of Birmingham (n. 2794224 Collaboration Agreement “ImmunoPEP”). ASav is supported by “TRAVEL (TRAnslation & deVELopment)” Scholarship (Department of Pharmacy, University of Naples Federico II, FARM/BF/02/2024). NM is supported by AORN A. Cardarelli Scholarship (n. 725/2022 GRC-Linea Progettuale 1.3: “Gestione delle cronicità”). ASch is supported by University of Naples Federico II PhD scholarship in “Nutraceuticals, functional foods and human health” (PNRR DM 118 M4C1– INV 4.1 ricerca PNRR generici). MS was funded by The Scientific and Technological Research Council of Türkiye (TÜBİTAK) 2214-A - International Research Fellowship Programme for PhD Students

## Ethics Approval Statement

Animal studies in Birmingham were regulated by the Animals (Scientific Procedures) Act 1986 of the United Kingdom and performed under Personal Project Licence P379E5607. Approval was granted by the University of Birmingham’s Animal Welfare and Ethical Review Body and all ethical guidelines were adhered to whilst carrying out this study. Blood was collected under approval from the University of Birmingham Local Ethical Review Committee (ERN_07-058). For studies in Italy (*in vivo* model of psoriasis), all animal care and experimental procedures complied with reporting of *in vivo* experiments (ARRIVE) guidelines, international and national law and policies and were approved (Authorisation number: 507/2022-PR) by the Italian Ministry of Health (EU Directive 2010/63/EU for animal experiments, and the Basel declaration including the 3Rs concept).

## Data availability

Data are available upon reasonable request. All data associated with this study are present in the paper or the online supplemental materials. Requests for reagents should be directed to the corresponding author and will be made available after completion of a material transfer agreement with *ImmunoPharmaLab*, Department of Pharmacy, University of Naples Federico II and/or the University of Birmingham.

## Contributions

ASav, BA, ST, LP, MS, AF, AM, MC, AJI, NM, Asch, SH, contributed to investigation and formal analysis. FM, MC, AJI, HMM AM, and GER contributed to conceptualisation, formal analysis, funding acquisition, project administration, resources, supervision and writing - original draft. All authors contributed to the writing, review and editing.

## References

1 Bayry, J & Radstake, T.R. Immune-mediated inflammatory diseases: progress in molecular pathogenesis and therapeutic strategies. Expert Rev. Clin. Immunol. 9, 297 (2013). 10.1586/eci.13.10

2 Atzeni, F. et al. Investigating the Potential Side Effects of Anti-TNF Therapy for Rheumatoid Arthritis: Cause for Concern? Immunotherapy. 7:353 (2015). doi: 10.2217/imt.15.4.

3 Taylor, P.C., Cerinic, M.M., Alten, R., Avouac, J., & Westhovens, R. Managing inadequate response to initial anti-TNF therapy in rheumatoid arthritis: optimising treatment outcomes. Ther Adv Musculoskelet Dis 14, 1 (2022). Doi 10.1177/1759720X221114101

4 Vik, A., Dalli, J., & Hansen JV. Recent advances in the chemistry and biology of anti-inflammatory and specialized pro-resolving mediators biosynthesized from n-3 docosapentaenoic acid. Bioorg Med Chem Lett. 27, 2259 (2017). 10.1016/j.bmcl.2017.03.079

5 Iyer, S.S., & Cheng G. Role of Interleukin 10 Transcriptional Regulation in Inflammation and Autoimmune Disease. Crit Rev Immunol.; 32, 23 (2012). doi: 10.1615/critrevimmunol.v32.i1.30

6 Chimen, M. et al. Homeostatic regulation of T cell trafficking by a B cell–derived peptide is impaired in autoimmune and chronic inflammatory disease. Nature Medicine., 21, 467 (2015). doi: 10.1038/nm.3842

7 Pezhman, L. et al. PEPITEM modulates leukocyte trafficking to reduce obesity-induced inflammation. Clin Exp Immunol 7, 212 (2023). doi: 10.1093/cei/uxad022.

8 Matsubara, H. et al. PEPITEM/Cadherin 15 Axis Inhibits T Lymphocyte Infiltration and Glomerulonephritis in a Mouse Model of Systemic Lupus Erythematosus. J Immunol 204 2043 (2020). 10.4049/jimmunol.1900213.

9 Hopkin, S.J., et al. Rejuvenation of leukocyte trafficking in aged mice through PEPITEM intervention. NPJ Aging 10, 33 (2024). doi: 10.1038/s41514-024-00160-6.

10 Kuna, M., Mahdi, F., Chade, A.R. & Bidwell G.L. Molecular Size Modulates Pharmacokinetics, Biodistribution, and Renal Deposition of the Drug Delivery Biopolymer Elastin-like Polypeptide. Sci Reports 8, 7923 (2018). DOI:10.1038/s41598-018-24897-9.

11 Deacon, C.F., Johnsen, A.H. & Holst, J.J. Degradation of glucagon-like peptide-1 by human plasma in vitro yields an N-terminally truncated peptide that is a major endogenous metabolite in vivo. J Clin Endocrin & Met 80, 952 (1995). 10.1210/jcem.80.3.7883856

12 Di L. Strategic Approaches to Optimizing Peptide ADME Properties. AAPS J 17, 134 (2015). doi: 10.1208/s12248-014-9687-3

13 Armstrong, A.W. & Read C. Pathophysiology, Clinical Presentation, and Treatment of Psoriasis: A Review. JAMA 323, 1945 (2020). doi:10.1001/jama.2020.4006.

14 Gauthier, T., Liu, D. (2022). Peptide Synthesis: Methods and Protocols. In: Jois, S.D. (eds) Peptide Therapeutics. AAPS Advances in the Pharmaceutical Sciences Series, vol 47. Springer, Cham. 10.1007/978-3-031-04544-8_2

15 Saviano, A. et al. Anti-inflammatory and immunomodulatory activity of Mangifera indica L. reveals the modulation of COX-2/mPGES-1 axis and Th17/Treg ratio. Pharmacol Res 182, 106283 (2022). doi: 10.1016/j.phrs.2022.106283

16 Saviano, A. et al. New biologic (Ab-IPL-IL-17) for IL-17-mediated diseases: identification of the bioactive sequence (nIL-17) for IL-17A/F function. Ann Rheum Dis 82, 1415 (2023). doi: 10.1136/ard-2023-224479. Epub 2023 Aug 14. PMID: 37580108; PMCID: PMC10579190.

17 Nguyen, L.T.H., Choi, M.J., Shin, H.M. & Yang, I.J. Coptisine Alleviates Imiquimod-Induced Psoriasis-like Skin Lesions and Anxiety-like Behaviour in Mice. Molecules 27, 1412 (2022). doi: 10.3390/molecules27041412. PMID: 35209199; PMCID: PMC8878104.

18 Chuo, W.H. et al. Alantolactone Suppresses Proliferation and the Inflammatory Response in Human HaCaT Keratinocytes and Ameliorates Imiquimod-Induced Skin Lesions in a Psoriasis-Like Mouse Model. Life (Basel) 11, 616 (2021). doi: 10.3390/life11070616. PMID: 34202301; PMCID: PMC8303865.

19 Gao, J. et al. Daphnetin inhibits proliferation and inflammatory response in human HaCaT keratinocytes and ameliorates imiquimod-induced psoriasis-like skin lesion in mice. Biol Res 53, 48 (2020). doi: 10.1186/s40659-020-00316-0. PMID: 33081840; PMCID: PMC7576854.

20 Raucci, F., ET AL. IL-17-induced inflammation modulates the mPGES-1/PPAR-γ pathway in monocytes/macrophages. Br J Pharmacol 179, 1857 (2022). doi: 10.1111/bph.15413. Epub 2021 Mar 21. PMID: 33595097.

21 Maione, F. et al. Interleukin 17 sustains rather than induces inflammation. Biochem Pharmacol 77, 878 (2009). doi: 10.1016/j.bcp.2008.11.011.

22 Saviano, A. Et al. A reverse translational approach reveals the protective roles of Mangifera indica in inflammatory bowel disease. J Autoimmun 144, 103181 (2024). doi: 10.1016/j.jaut.2024.103181.

23 Mangan, P.R. et al. Dual Inhibition of Interleukin-23 and Interleukin-17 Offers Superior Efficacy in Mouse Models of Autoimmunity. J Pharmacol Exp Ther 354, 152 (2015) doi: 10.1124/jpet.115.224246. Epub 2015 May 26. PMID: 26015463.

24 Moos, S., Mohebiany, A.N., Waisman, A. & Kurschus, F.C. Imiquimod-Induced Psoriasis in Mice Depends on the IL-17 Signaling of Keratinocytes. J Invest Dermatol 139, 1110 (2015). doi: 10.1016/j.jid.2019.01.006.

25 Wang, D. et al. Calcium/calmodulin-dependent protein kinase IV promotes imiquimod-induced psoriatic inflammation via macrophages and keratinocytes in mice. Nat Commun. 13, 4255 (2022). doi: 10.1038/s41467-022-31935-8.

26 Ippagunta, S.K. et al. Keratinocytes contribute intrinsically to psoriasis upon loss of Tnip1 function. Proc Natl Acad Sci USA 113, E6162 (2016). doi: 10.1073/pnas.1606996113.

27 Yong, L. et al. Targeting the transcription factor HES1 by L-menthol restores protein phosphatase 6 in keratinocytes in models of psoriasis. Nat Commun 13, 7815 (2022). doi: 10.1038/s41467-022-35565-y.

28 van der Fits, L. et al. Imiquimod-induced psoriasis-like skin inflammation in mice is mediated via the IL-23/IL-17 axis. J Immunol 182, 5836 (2009). doi: 10.4049/jimmunol.0802999. PMID: 19380832.

29 Horváth, S. et al, Methodological refinement of Aldara-induced psoriasiform dermatitis model in mice. Sci Rep 9, 3685 (2019). doi: 10.1038/s41598-019-39903-x. PMID: 30842501; PMCID: PMC6403245.

30 Ha, H.L. et al. IL-17 drives psoriatic inflammation via distinct, target cell-specific mechanisms. Proc Natl Acad Sci USA 111, E3422 (2014). doi: 10.1073/pnas.1400513111.

31 Lou, F., Sun, Y. & Wang, H. Protocol for Flow Cytometric Detection of Immune Cell Infiltration in the Epidermis and Dermis of a Psoriasis Mouse Model. STAR Protoc 1, 100115 (2020). doi: 10.1016/j.xpro.2020.100115. PMID: 33377011; PMCID: PMC7757015.

32 Regan-Komito, D. et al. Absence of the Non-Signalling Chemerin Receptor CCRL2 Exacerbates Acute Inflammatory Responses In Vivo. Front Immunol 8, 1621 (2018). doi: 10.3389/fimmu.2017.01621.

33 Cossarizza, A. et al. Guidelines for the use of flow cytometry and cell sorting in immunological studies (third edition). Eur J Immunol 51, 2708 (2021). doi: 10.1002/eji.202170126.

34 Cristiano, C. et al. Neutralization of IL-17 rescues amyloid-β-induced neuroinflammation and memory impairment. Br J Pharmacol 176, 3544 (2019). doi: 10.1111/bph.14586. Epub 2019 Mar 3. PMID: 30673121; PMCID: PMC6715610.

35 Maione, F. et al. Analysis of the inflammatory response in HY-TCR transgenic mice highlights the pathogenic potential of CD4-CD8-T cells. Autoimmunity 43, 672 (2010). doi: 10.3109/08916931003678296.

36 Ercolano, G. et al. PPARɣ drives IL-33-dependent ILC2 pro-tumoral functions. Nat Commun 12, 2538 (2021). doi: 10.1038/s41467-021-22764-2. PMID: 33953160; PMCID: PMC8100153.

37 Saviano, A. et al. Supplementation with ribonucleotide-based ingredient (Ribodiet®) lessens oxidative stress, brain inflammation, and amyloid pathology in a murine model of Alzheimer. Biomed Pharmacother. 139, 111579 (2021). doi: 10.1016/j.biopha.2021.111579.

38 Saviano, A., et al. In Silico, In Vitro, and In Vivo Analysis of Tanshinone IIA and Cryptotanshinone from Salvia miltiorrhiza as Modulators of Cyclooxygenase-2/mPGES-1/Endothelial Prostaglandin EP3 Pathway. Biomolecules. 12, 99 (2022). doi: 10.3390/biom12010099. PMID: 35053247; PMCID: PMC8774285.

39 Caso, F. et al. Analysis of rheumatoid- vs psoriatic arthritis synovial fluid reveals differential macrophage (CCR2) and T helper subsets (STAT3/4 and FOXP3) activation. Autoimmun Rev 21, 103207 (2022). doi: 10.1016/j.autrev.2022.103207. Epub 2022 Oct 1. PMID: 36191778.

40 Vellecco, V. et al. Interleukin-17 (IL-17) triggers systemic inflammation, peripheral vascular dysfunction, and related prothrombotic state in a mouse model of Alzheimer’s disease. Pharmacol Res. 187, 106595 (2023). doi: 10.1016/j.phrs.2022.106595.

41 Di Meglio, P., Perera, G.K. & Nestle, F.O. The multitasking organ: recent insights into skin immune function. Immunity 356, 857 (2011). doi: 10.1016/j.immuni.2011.12.003.

42 Perera, G.K., Di Meglio, P. & Nestle, F.O. Psoriasis. Annu Rev Pathol 7, 385 (2012). doi: 10.1146/annurev-pathol-011811-132448.

43 van der Fits, L. et al. Imiquimod-Induced Psoriasis-Like Skin Inflammation in Mice Is Mediated via the IL-23/IL-17 Axis1. J Immunol 182, 5836 (2009).

44 Pender, E.K. & Kirby, B. An update on topical therapies for psoriasis. Curr Opin Rheumatol 36, 289 (2024). doi: 10.1097/BOR.0000000000001018.

45 Lewis J.W., et al. Therapeutic avenues in bone repair: Harnessing an anabolic osteopeptide, PEPITEM, to boost bone growth and prevent bone loss. Cell Rep Med. 2024 May 21;5(5):101574.doi: 10.1016/j.xcrm.2024.101574.

