## Supplementary Tables and Figures for "PEPITEM Tripeptides and Peptidomimetics: Next-Generation Modulators of Inflammation in Immune-Mediated Conditions"

**Supplementary Table 1. The sequence of peptides used in the alanine sweep of the PEPITEM molecule.**

| Peptide number | |
|---|---|
| 1 | AVTEQGAELSNEER |
| 2 | SATEQGAELSNEER |
| 3 | SVAEQGAELSNEER |
| 4 | SVTAQGAELSNEER |
| 5 | SVTEAGAELSNEER |
| 6 | SVTEQAAELSNEER |
| 7 (PEPITEM) | SVTEQGAELSNEER |
| 8 | SVTEQGAALSNEER |
| 9 | SVTEQGAEASNEER |
| 10 | SVTEQGAELANEER |
| 11 | SVTEQGAELSAEER |
| 12 | SVTEQGAELSNAER |
| 13 | SVTEQGAELSNEAR |
| 14 | SVTEQGAELSNEEA |

**Supplementary Table 2 The physical and pharmacological characteristics of PEPITEM, its pharmacophores and its peptidomimetics.**

| Sequence | MW | % inhibition of Lymphocyte trafficking (0.3nM) | $IC_{50}$ (pM) | T½ in plasma (min; max =180min) | Designation | Structure |
|---|---|---|---|---|---|---|
| PEPITEM | 1549 | 40 | 45 | >180 | Native sequence | 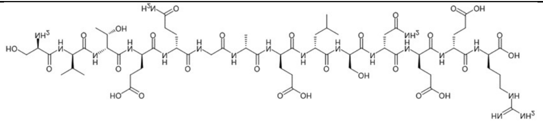 |
| SVT-acid | 305 | 42 | 46 | 20 | Native sequence | 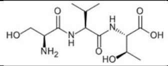 |
| Ac-QGA-acid | 316 | 38 | 42 | ND | Native sequence | 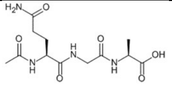 |
| [PyroGlu]-GA-NH2 | 256 | 55 | 3.2 | >180 | **Peptidomimetics ↓** | 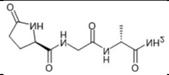 |
| SVT-NH-ethyl | 332 | 46 | 17 | >180 | | 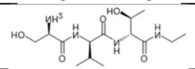 |
| tvs-NH2 | 304 | 59 | 24 | >180 | | 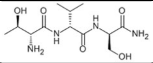 |
| ***SVT based series ↓*** | | | | | | |
| Ac-SVT-acid | 347 | 37 | ND | ND | | 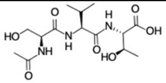 |
| SVT-NH2 | 304 | 39 | 10 | >180 | | 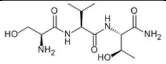 |
| TVT-NH2 | 318 | 45 | ND | 17 | | 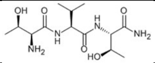 |
| SVS-NH2 | 290 | 43 | ND | 103 | | 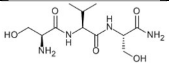 |
| SLT-NH2 | 318 | 46 | ND | 7 | | 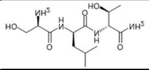 |
| SWT-acid | 391 | 47 | ND | 74 | | 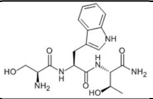 |
| SYT-acid | 368 | 39 | ND | 17 | | 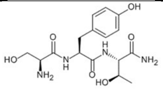 |

| | | | | | | | |
|---|---|---|---|---|---|---|---|
| S(Me)-VT-NH2 | 318 | 35 | ND | ND | | | 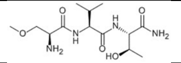 |
| sVT-NH2 | 304 | 9 | ND | ND | | | 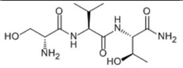 |
| svt-NH2 | 304 | 25 | ND | ND | | | 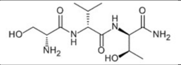 |
| **QGA based series ↓** | | | | | | | |
| Ac-QGA-NH2 | 315 | 40 | 100 | >180 | | | 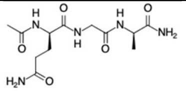 |
| Ac-QGA-NH-ethyl | 343 | 20 | ND | ND | | | 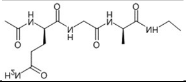 |
| AcNGA-NH2 | 301 | 32 | ND | ND | | | 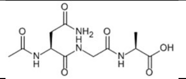 |
| [PyroGlu]-GA-NH-ethyl | 284 | 35 | ND | >180 | | | 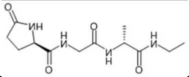 |
| [PyroGlu]-GA-NH-methyl | 270 | 40 | ND | 97 | | | 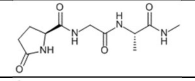 |
| [PyroGlu]-GL-NH2 | 298 | 25 | ND | >180 | | | 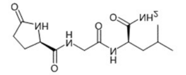 |
| [PyroGlu]-GF-NH2 | 332 | 49 | ND | 146 | | | 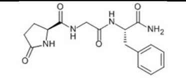 |
| [D-pyroGlu]-Ga-NH2 | 256 | 46 | ND | 70 | | | 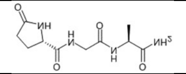 |
| Ac-qGa-NH2 | 315 | 34 | ND | ND | | | 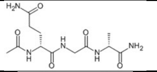 |
| aGq-acid | 274 | 26 | ND | ND | | | 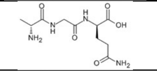 |
| Ac-QG[Aib]-acid | 330 | 32 | ND | ND | | | 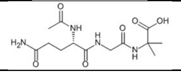 |

The molecular weight (MW), % inhibition of lymphocyte trafficking at a concentration of 0.3nM, IC50 (ND = Not done), half-life in plasma, designation (Native sequence or peptidomimetic), chemical structure of PEPITEM, its pharmacophores and its peptidomimetics.

**Supplementary Table 3. The calculated physicochemical properties of PEPITEM its pharmacophores and peptidomimetics**

| AA Seq | Number AA | MW | logD | logP | TPSA | Rotatable Bonds | HBD | HBA |
|---|---|---|---|---|---|---|---|---|
| SVTEQGAELSNEER | 14 | 1549 | 3.083 | -4.68 | 799.6 | 68 | 27 | 47 |
| SVT-acid | 3 | 305.3 | -2.679 | -3.605 | 162 | 10 | 6 | 9 |
| Ac-QGA-acid | 3 | 316.3 | -1.8 | -2.278 | 167.7 | 12 | 5 | 10 |
| [PyroGlu]-GA-NH2 | 3 | 256.3 | -2.401 | -2.401 | 130.4 | 7 | 4 | 8 |
| SVT-NH-Ethyl | 3 | 332.4 | -0.7978 | -1.442 | 153.8 | 12 | 6 | 9 |
| tvs-NH2 | 3 | 304.3 | -1.41 | -2.026 | 167.8 | 10 | 6 | 9 |
| Ac-SVT-acid | 3 | 347.4 | -1.456 | -1.397 | 165.1 | 12 | 6 | 10 |
| SVT-NH2 | 3 | 304.3 | -1.366 | -2.026 | 167.8 | 10 | 6 | 9 |
| TVT-NH2 | 3 | 316.4 | -0.4568 | -0.8411 | 147.5 | 10 | 5 | 8 |
| SVS-NH2 | 3 | 290.3 | -1.597 | -2.399 | 167.8 | 10 | 6 | 9 |
| SLT-NH2 | 3 | 304.3 | -1.455 | -2.06 | 167.8 | 11 | 6 | 9 |
| SWT-acid | 3 | 378.4 | -2.053 | -2.942 | 177.8 | 11 | 7 | 10 |
| SYT-acid | 3 | 354.4 | -1.791 | -2.268 | 188 | 11 | 7 | 10 |
| S(Me)-VT-NH2 | 3 | 318.4 | -1.048 | -1.843 | 156.8 | 11 | 5 | 9 |
| sVT-NH2 | 3 | 304.3 | -1.366 | -2.026 | 167.8 | 10 | 6 | 9 |
| svt-NH2 | 3 | 304.3 | -1.366 | -2.026 | 167.8 | 10 | 6 | 9 |
| Ac-QGA-NH2 | 3 | 315.3 | -2.629 | -2.629 | 173.5 | 12 | 5 | 10 |
| Ac-QGA-NH-Ethyl | 3 | 343.4 | -1.805 | -1.805 | 159.5 | 14 | 5 | 10 |
| Ac-NGA-NH2 | 3 | 301.3 | -2.657 | -2.657 | 173.5 | 11 | 5 | 10 |
| [PyroGlu]-GA-NH-Ethyl | 3 | 284.3 | -1.513 | -1.513 | 116.4 | 9 | 4 | 8 |
| [PyroGlu]-GA-NH-Methyl | 3 | 270.3 | -1.959 | -1.959 | 116.4 | 8 | 4 | 8 |
| [PyroGlu]-GL-NH2 | 3 | 298.3 | -1.34 | -1.34 | 130.4 | 9 | 4 | 8 |
| [PyroGlu]-GF-NH2 | 3 | 332.4 | -1.068 | -1.068 | 130.4 | 9 | 4 | 8 |
| [D-pyroGlu]-Ga-NH2 | 3 | 256.3 | -2.401 | -2.401 | 130.4 | 7 | 4 | 8 |
| Ac-qGa-NH2 | 3 | 315.3 | -2.629 | -2.629 | 173.5 | 12 | 5 | 10 |
| aGq-acid | 3 | 274.3 | -2.782 | -4.128 | 164.6 | 10 | 5 | 9 |
| Ac-QG[Aib]-acid | 3 | 330.3 | -1.678 | -1.923 | 167.7 | 12 | 5 | 10 |

Physicochemical properties calculated using Optibrium StarDrop version 7.6.1 software

**CERTIFICATE OF ANALYSIS** BIOSYNTH

| | |
|---|---|
| *Cat No.* | PEP-00023262 |
| *Product* | H-SVTEQGAELSNEER-OH |

| | | | |
|---|---|---|---|
| *CAS RN* | n.a. | *M. F.* | n.a. |
| *Lot No.* | SX52-A0306 | *M. W.* | 1548.58 g/mol |
| | | *Salt Form* | Trifluoroacetate (TFA) Salt |

| *Test* | *Result* | *Specification* |
|---|---|---|
| Purity (UPLC 215 nm) | 97.2 % | min. 95.0 % |
| Identification (ESI MS) | 1547.6 | theoretical monoisotopic molecular weight MW: 1547.7 ± 1 u |

*Date of issue:* May 30, 2024 — BIOSYNTH Group

*PA No.:* 141013 — Anan Mohammed

*Internal Reference No.:* 4500138980

154667 - V08

*Product: Research Grade Custom Peptide*
*Notes applicable:*

*Storage: Refer to the Quality Control Detail Information on our website at https://www.biosynth.com/uploads/FAQ/peptide-storage-tips.pdf*

Sequence: H-SVTEQGAELSNEER-OH — 29/May/2024
Customer Code: SX52-A0306 — Column: ACE 3 C18-300 150x2,1mm — Flow: 0,35mL/min — 2-70% 215 nm,amx
MW: 1548,58 — Eluent: A_H2O+0.05%TFA ; B_ACN+0.05%TFA

Sample Name SX52-A0306_UV

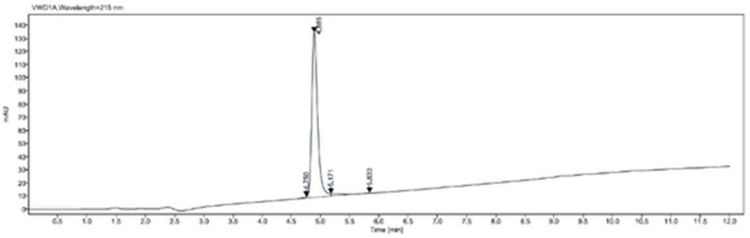


Signal: VWD1A,Wavelength=215 nm

| RT [min] | Height | Area | Area% |
|---|---|---|---|
| 4.750 | 1.0583 | 3.7368 | 0.4159 |
| 4.885 | 125.8811 | 873.0886 | 97.1840 |
| 5.171 | 1.7223 | 19.1955 | 2.1367 |
| 5.833 | 0.2990 | 2.3665 | 0.2634 |
| | Sum | 898.3874 | |

Spectrum Plot Report — Agilent

| | | | | | | | |
|---|---|---|---|---|---|---|---|
| Name | SX52-A0306_mass | Rack Pos. | | Instrument | Instrument 1 | Operator | |
| Inj. Vol. (ul) | 25 | Plate Pos. | | IRM Status | All ions missed | | |
| Data File | SX52-A0306_mass.d | Method (Acq) | ACE_Fast FINAL.m | Comment | | Acq. Time (Local) | 29/05/2024 16:05:00 (UTC+01:00) |

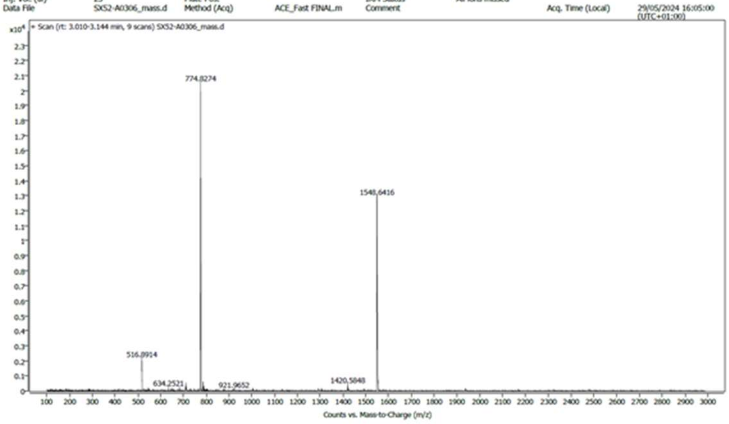


**Supplementary Fig. 1.** Certificate of analysis and purification reports for PEPITEM (SVTEQGAELSNEER) form the CRO Biosynth.

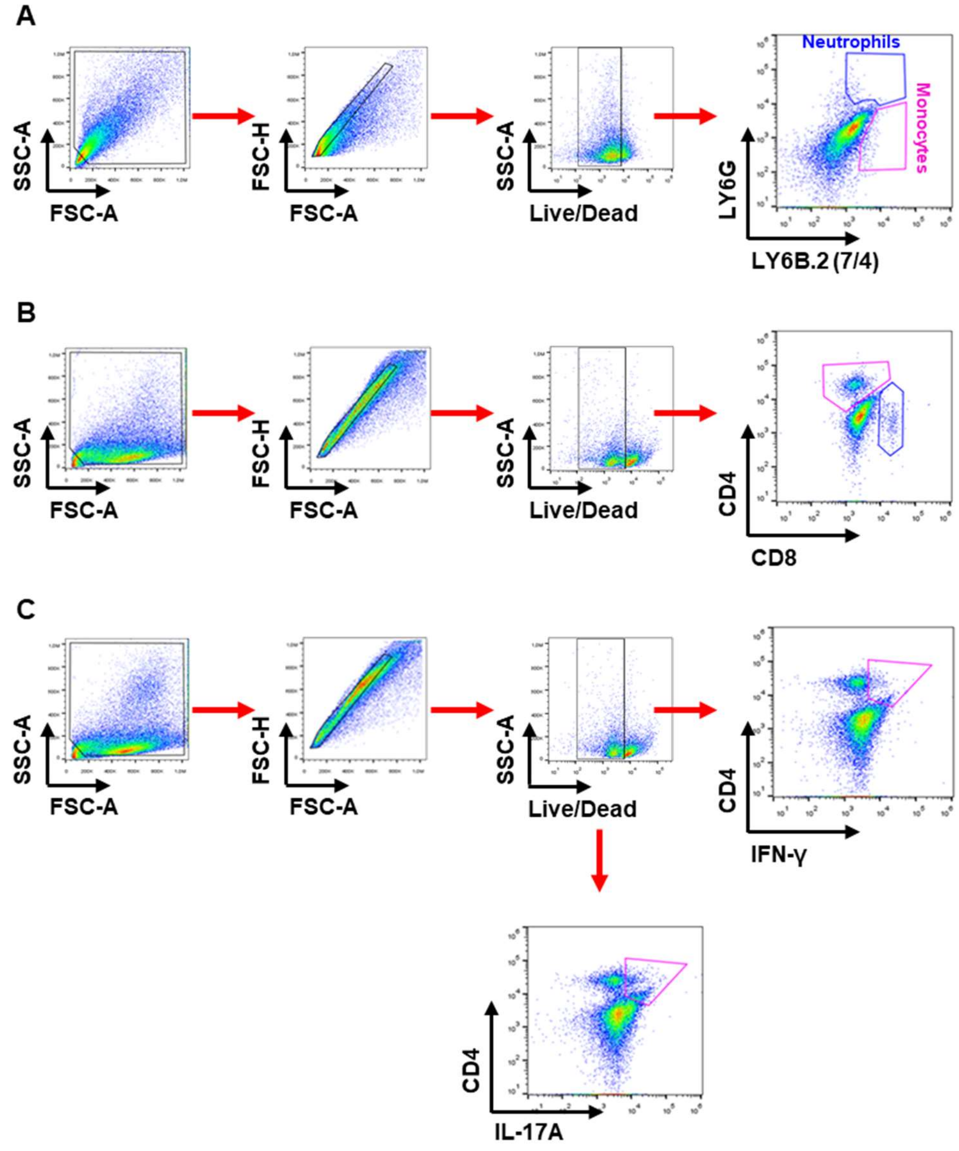


**Supplementary Fig. 2. A)** Flow cytometry strategy applied to identify skin infiltrated neutrophils and monocytes and **B) and C)** splenic lymphocyte populations. FACS pictures are presented as dot plot (pseudocolor).

.

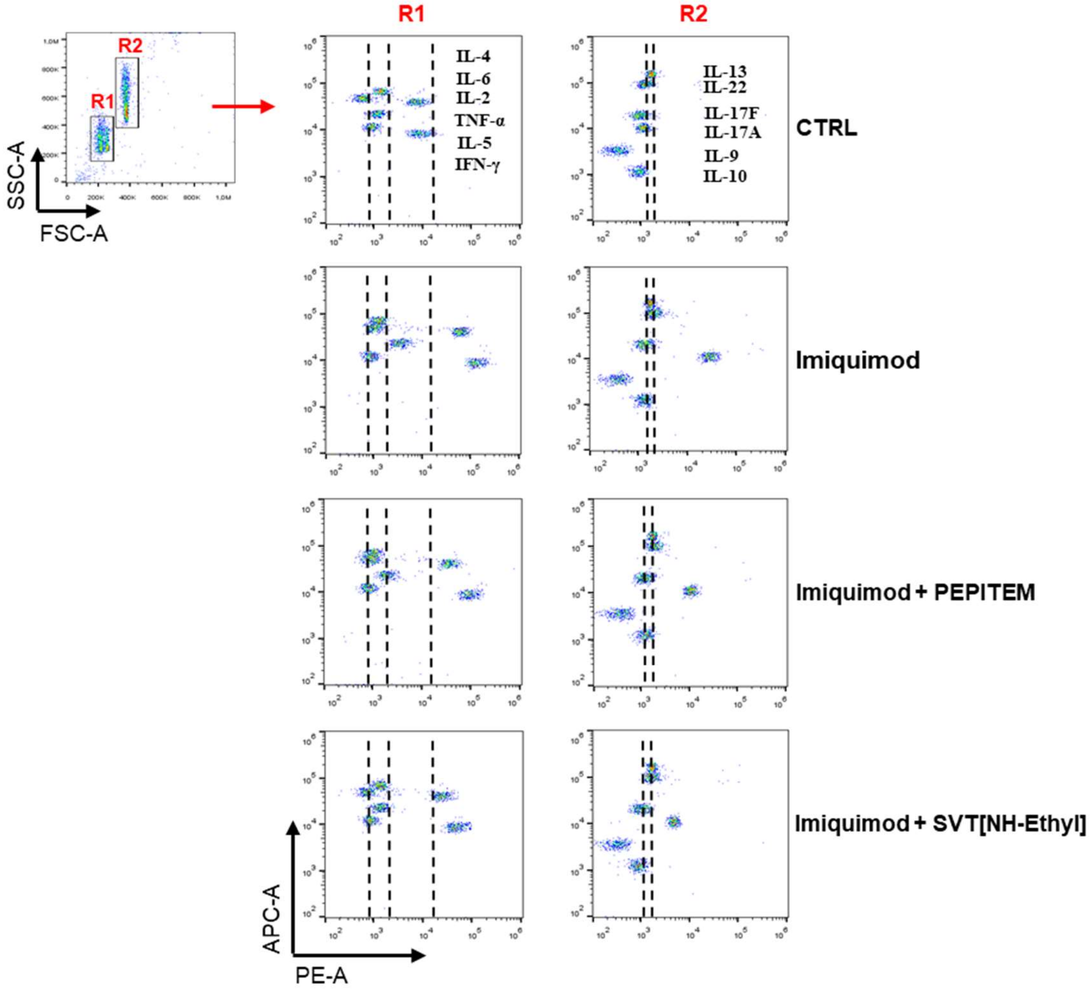


**Supplementary Fig. 3.** Flow cytometry gating strategy for multi-LEGENDplex™ Th-related cytokine assay.

A

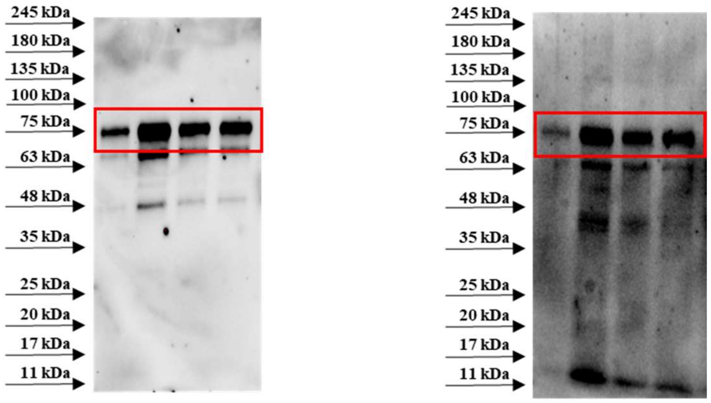


B

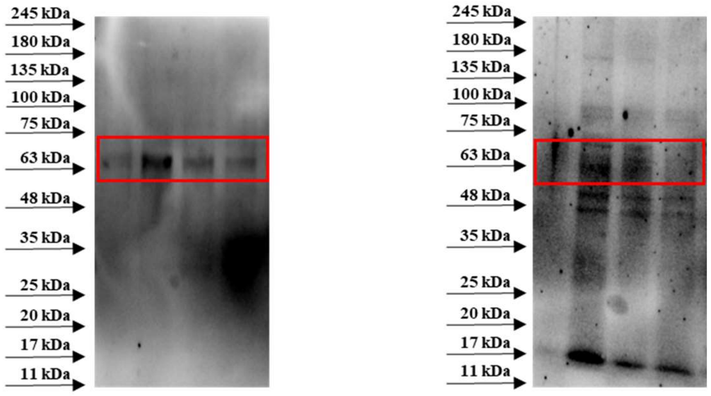


C

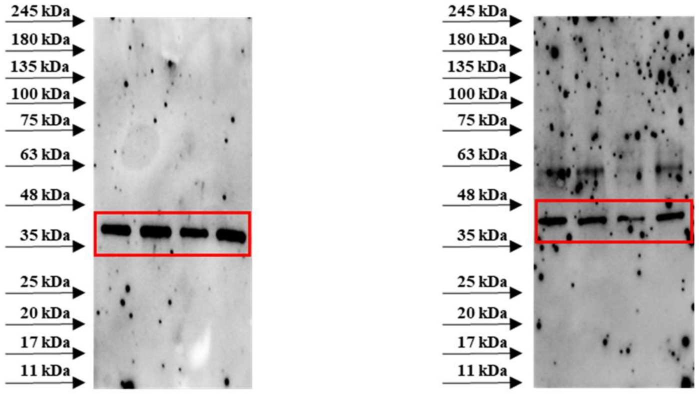


**Supplementary Fig. 4.** Uncropped western blots for **A)** COX-2 (~72 kDa), **B)** NF-κB-p65 (~65 kDa) and **C)** β-actin (~42 kDa) obtained from skin lesions homogenates in all experimental conditions (CTRL, Imiquimod, Imiquimod + PEPITEM, Imiquimod + SVT[NH-Ethyl], from left to right respectively). Images represent technical duplicate of one experiment run with n=6 mice per group pooled.

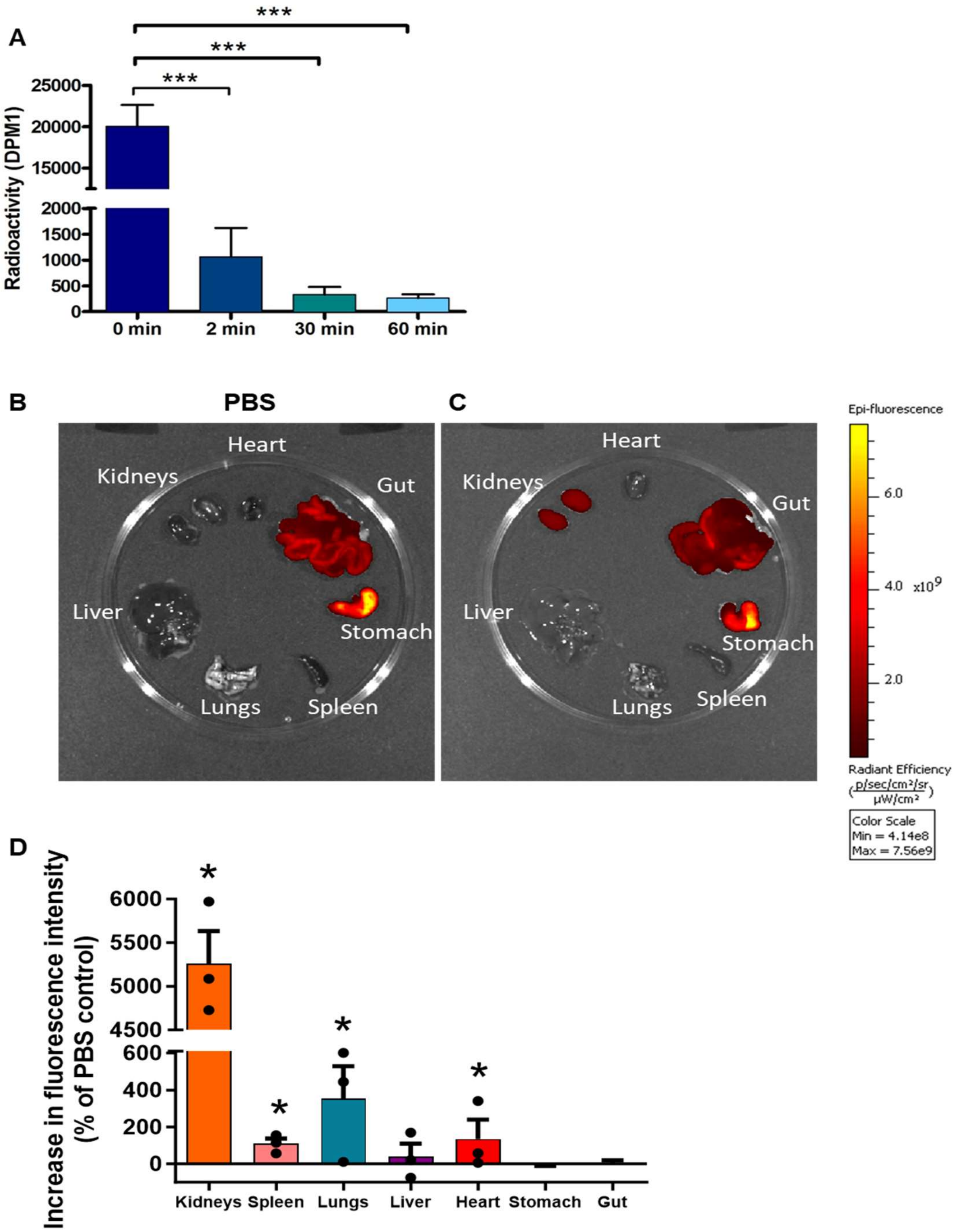


**Supplementary Fig. 5.** The pharmacokinetics of PEPITEM *in vivo*. **A)** 1ng/g $^{3}H$ PEPITEM was injected via the tail vein of C57BL/6 mice for 2-60 min. Blood was extracted and assessed for radioactivity by scintillation counting. Radioactivity at 0 min was determined by spiking 1ng/g $^{3}H$ into collected blood before centrifugation and measurement of radioactivity. Data expressed as radioactive counts (DPM1), and represents mean ± SEM of 3-6 mice. $P \leq 0.001$ by one way ANOVA followed by Dunnett's post-test. **B) and C)** IVIS imaging of organs following IV injection of PBS or PEPITEM-AF680 into BALB/C mice. Organs were imaged *ex-vivo* after 15 min. Representative images taken using the IVIS Spectrum Imager, with 3 mice per group. Fluorescence is expressed radiant efficiency (p/s/cm$^{2}$/sr). **D)** The organ distribution of PEPITEM-AF680 *in vivo*. Data are mean ± SEM of 3 experiments expressed as % of PBS control. * = $P \leq 0.05$ by Mann-Whitney t-test. N=3 mice per group. Data expressed as mean ± SEM. N=3.

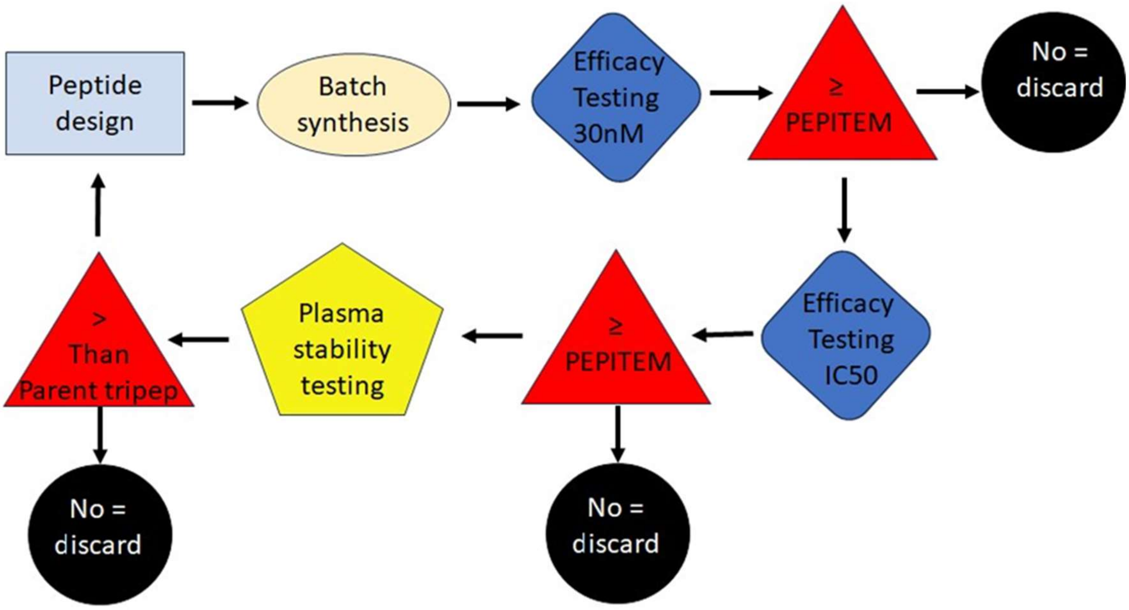


**Supplementary Fig. 6.** The strategy for iterative peptide optimisation of the tripeptides SVT and QGA.

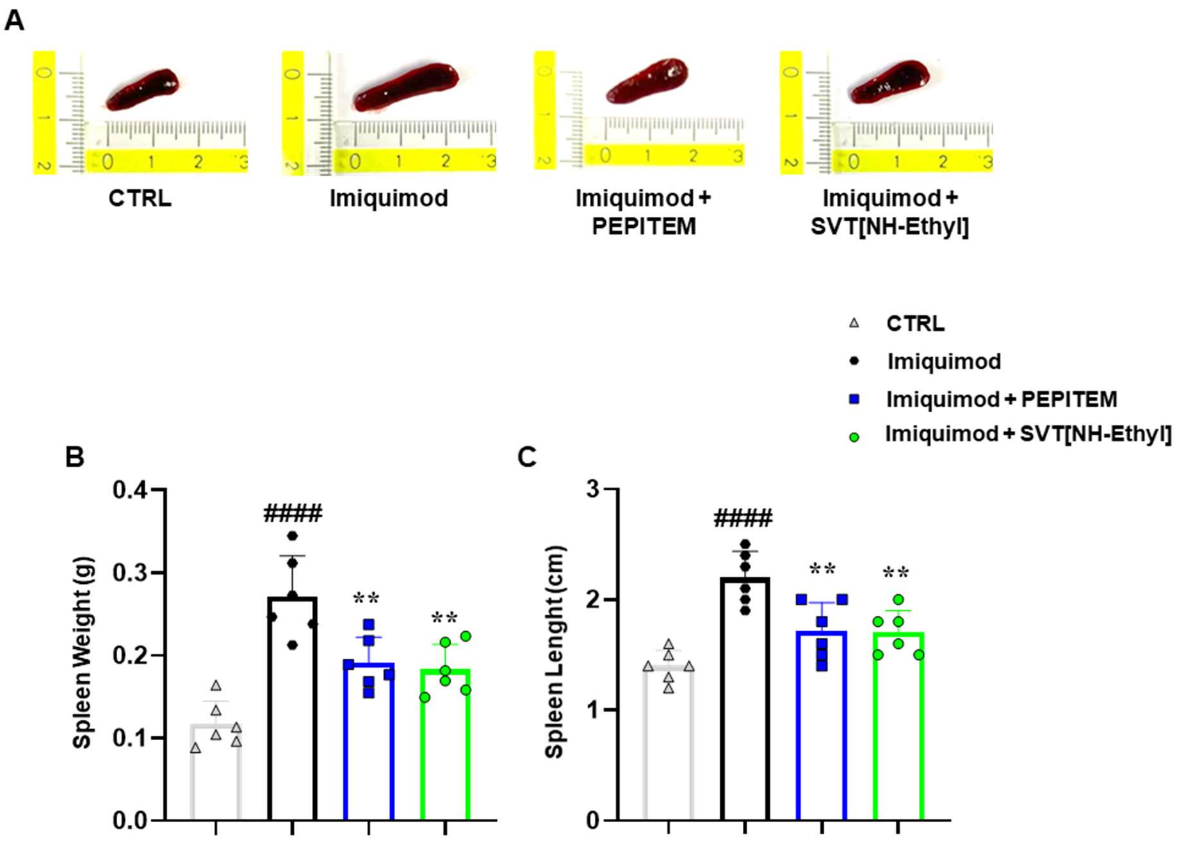


**Supplementary Fig. 7.** At the experimental endpoint (day 7) spleen from CTRL, Imiquimod, Imiquimod + PEPITEM and Imiquimod + SVT[NH-Ethyl] groups were dissected and analysed macroscopically. **A)** Representative photographs of organ necroscopy, **B)** weight (indicated as g) and **C)** length (indicated as cm) were evaluated. Data are presented as means ± S.D. of N=6 mice per group. Statistical analysis was conducted by one-way ANOVA followed by Bonferroni's for multiple comparisons. $^{####}$P < 0.0001 *vs* CTRL group; $^{**}$P < 0.01, *vs* Imiquimod group.
